# A microRNA-based Therapy against Human Medulloblastoma Validated in a Xenotransplant Model using Wild-Type Mouse Embryos

**DOI:** 10.1101/2025.04.25.650572

**Authors:** Carlotta Spattini, Letizia La Rosa, Rui C. Pereira, Lidia Giantomasi, Roberta Pelizzoli, Kiril Tuntevski, Giuseppe Battaglia, Andrea Di Fonzo, Andrea Armirotti, Davide De Pietri Tonelli

## Abstract

Medulloblastoma (MB) is the most prevalent malignant pediatric brain tumor. Its poor prognosis is driven by therapy-resistant cancer stem-like cells (CSCs), which are challenging to study in traditional preclinical models. MicroRNAs (miRNAs) modulate tumor-related pathways by repressing oncogenic gene expression, a function lost in cancer. We assessed the therapeutic potential of miRNA restoration in human MB from distinct subgroups, validated in a xenotransplant model using wild-type (WT) mouse embryos. Multi-omics, imaging, and *in silico* analyses were employed to identify underlying mechanisms and therapeutic targets. Human MB cells formed tumors in embryonic mouse brains that mirrored key MB features, including vascularized CSC niches and metastases. Restoration of underexpressed miRNAs reduced tumor growth and invasiveness *in vitro* and *in vivo.* These miRNAs synergistically repressed gene networks related to cell adhesion and RNA metabolism, shared oncogenic pathways across MB subgroups. Our embryonic xenotransplant model provides a clinically relevant platform for evaluating next-generation gene therapies in pediatric brain cancer.

## INTRODUCTION

Medulloblastoma (MB) is one of the most common malignant pediatric brain tumors (World Health Organization –WHO– grade IV), with recent epidemiological studies indicating a significant rise in its prevalence^1^. Although MB prognosis has recently improved due to advancements in treatment, long-term survivors often experience severe side effects from radiation and cytotoxic chemotherapy, and mortality remains considerable^2^. A major challenge in MB treatment is tumor relapse, which occurs in approximately 30% of cases, with incidence rates varying among molecular subgroups^3^. MB relapse is largely attributed to the presence of cancer stem-like cell (CSC) subpopulations within the tumor, which can generate therapy-resistant clones following first-line treatment^4^. This phenomenon is especially pronounced in the Sonic Hedgehog (SHH) and Group 3, the most common and aggressive of the four MB subtypes, respectively (the other two MB groups being the Wingless-activated –WNT– and Group 4^4^. Furthermore, MB frequently presents with metastases complicating surgical resection and worsening the prognosis, particularly in groups 3 and 4^4^. These challenges highlight the urgent need for effective therapies that limit CSC expansion and tumor invasiveness^5^. Finally, MB arises from a heterogeneous landscape of mutations, spanning both coding and noncoding regions of the genome^6,7^.

Preclinical research, however, is complicated because the neural stem/precursors (NSC/NPCs) at the origin of this pediatric tumor are highly heterogeneous^8^, and their fate is regulated in a human-specific manner^9^. The current animal models fail to fully recapitulate the human embryonic cell types and signaling underlying MB initiation and progression^10^. This underscores the need for better alternative strategies to recapitulate the tumor developmental milieu, enabling more accurate therapeutic testing.

Due to this complexity, single-target therapies are unlikely to provide long-term efficacy against MB. In this context, microRNAs (miRNAs) — small noncoding RNAs that post-transcriptionally regulate gene expression — emerge as key players in MB initiation and progression^11^. Since the first evidence of miRNA dysregulation in cancer^12^, different studies have confirmed that altered miRNA abundance characterizes MB^13^. Many of the altered miRNAs in MB are essential for neural development^14^. This suggests that miRNA deregulation could be a causal factor in MB initiation and stemness^11^, making these molecules attractive therapeutic targets. Despite that, clinical trials targeting single miRNAs have largely failed in oncology, highlighting the need for alternative approaches^15^. Recent studies in other brain cancers, like glioblastoma (GBM), have demonstrated that combinatorial approaches, where multiple miRNAs are targeted simultaneously, enhance the efficacy of tumor suppression^16^. This aligns with the functional redundancy (*i.e.,* synergism) of miRNAs observed in neural development^14^. Yet, combinatorial miRNA therapy has not been tested in human MB.

This study validated a combination of miRNAs (henceforth “the Pool”) as a therapeutic strategy to reduce human MB growth and invasiveness. To assess the efficacy and stability of the Pool *in vivo*, we introduced a xenotransplant model using wild-type (WT) mouse embryos that recapitulates MB formation and progression in the context of a developing nervous system. Finally, through multi-omics analyses, we inferred mechanisms underlying the tumor-suppressive effects of these therapeutic miRNAs in cells from different human MB groups.

## RESULTS

### WT embryonic mouse brain supports engraftment and proliferation of human MB tumor

To implement an *in vivo* model suitable for the investigation of human MB biology in the context of neural development, we took advantage of the *in utero* orthotopic xenotransplantation strategy in mouse embryos, which we previously demonstrated to support the engraftment of human Glioblastoma and Meningioma, in WT^17,18^. Similarly to the models described above, we implemented the orthotopic xenotransplant of DAOY cells, one of the best characterized human cell lines of the SHH group for this cancer^19^. To facilitate the identification of human MB cells engrafted in the host, DAOY cells were transduced with a lentivirus expressing the mCherry fluorescent protein under the control of the constitutive elongation factor 1-alpha 1 (eEF1a1) promoter prior to transplantation. Then, a single-cell suspension of 50,000 DAOY mCherry^+^ cells was injected *in utero* into the lateral ventricle (LV) of WT mouse telencephalon at embryonic day 14.5 *post coitum* (E14.5) (Figure 1A). Virtually all the embryos that underwent mCherry^+^ DAOY cells xenotransplantation survived the procedure, an efficiency in line with previous reports^17,18^. Starting from E18.5, the injected brains were harvested and analyzed for the presence of MB tumor xenografts (Tx, Figure 1B-E). Previous studies reported fusion events between transplanted human embryonic NSC and host cells, generating neurons with hybrid mouse-human phenotypes^20^. Brain slices with mCherry^+^ Tx at E18.5 were examined by immunofluorescence staining for Human-specific nuclei (HuNu) antigen to assess this possibility in our model. This analysis revealed that all the DAOY mCherry^+^ cells were also co-stained for the HuNu marker (Figure 1B, *white arrows*), indicating that human MB cells retained their human identity once integrated into the mouse tissue, in line with previous reports^17,18^. Moreover, in agreement with studies demonstrating that species-specific morphological features of human nuclei can distinguish them from mouse cells^21^, nuclei of HuNu^+^/mCherry^+^ cells presented fewer nucleoli compared to nearby HuNu/mCherry negative cells (Figure 1B, white *arrowheads*). These results indicate that the WT embryonic mouse brain supports the engraftment of human MB cells.

**Figure 1.**
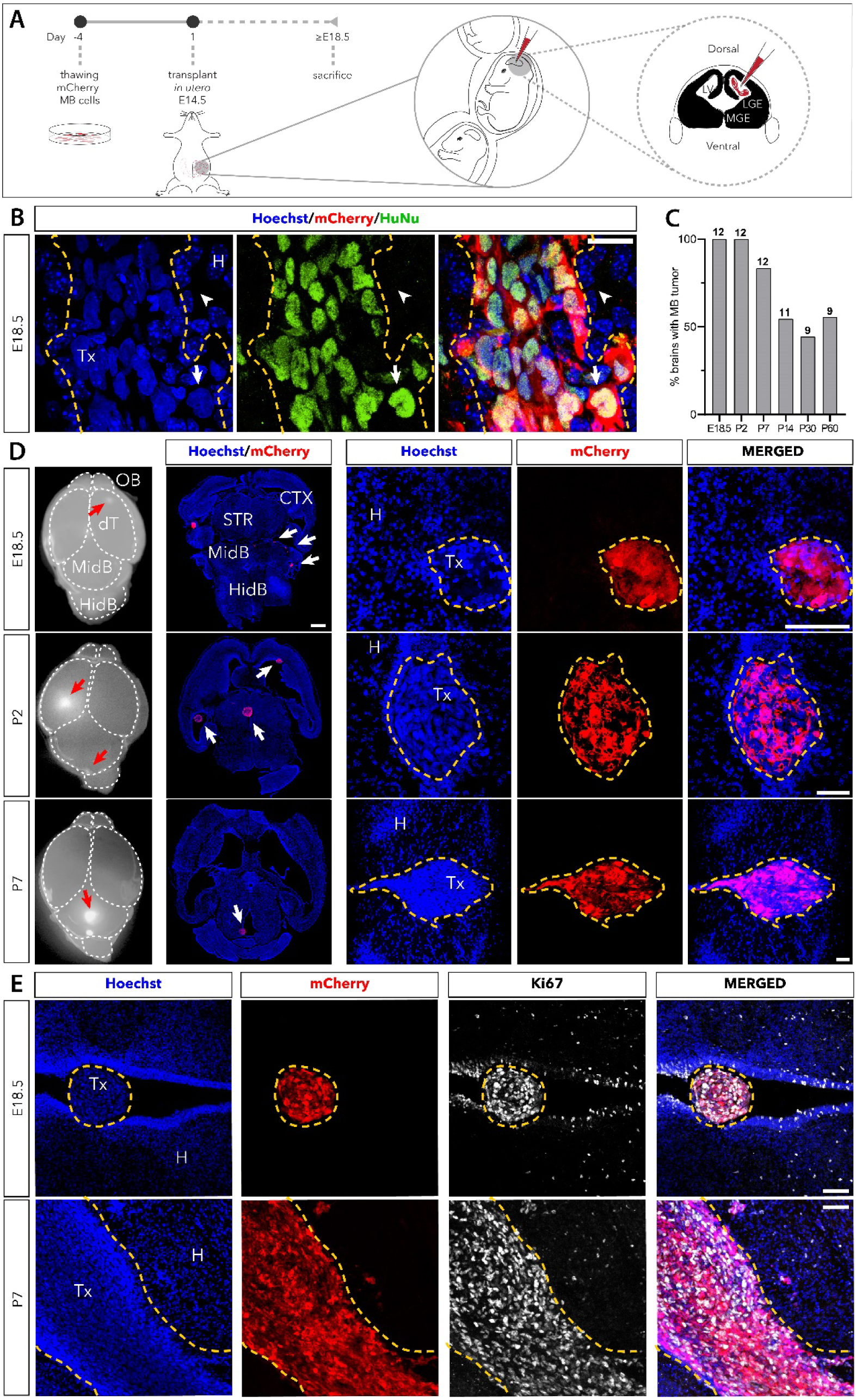
WT embryonic mouse brain supports engraftment and proliferation of human MB tumor. (A) Scheme of the experimental procedure: human MB cells were plated 5 days before the *in utero* transplantation and injected in the lateral ventricle (LV) of E14.5 embryos. (B) Immunofluorescence images of cryosections through the brain of E18.5 mouse embryo. Nuclei (Hoechst) of mCherry^+^ cells overlap with HuNu signal in the Tumor Xenograft (Tx, dashed line); arrows and arrowheads indicate human and mouse nuclei, respectively. Scale bar: 20µm. (C) Percentage of brains with Tx, over the total injected brains. Sample size (n= brains injected with DAOY cells) is reported at the top of each histogram bar. (D) Whole brains in dorsal view at the indicated time points; red arrows indicate the Tx (left); immunofluorescence images of cryosections through the brains at the indicated time points, showing mCherry^+^ Tx (white arrows) at low magnification, or at high magnification (right). Scale bar: 500µm. (E) Immunofluorescence images of cryosections at the indicated time points, showing Ki67^+^ cells within the Tx. Scale bar: 50µm. (H) Host tissue; Lateral Ventricle (LV); Lateral Ganglionic Eminence (LGE); Medial Ganglionic Eminence (MGE); Olfactory Bulb (OB); Dorsal Thalamus (dT); Midbrain (MidB); Hindbrain (HiB); and Striatum (STR).

At embryonic day 18.5 (E18.5) and postnatal day 2 (P2), all injected brains exhibited tumor engraftment (Tx; Figure 1C). This observation suggests that the developing immune system of wild-type (WT) mice tolerates the engraftment of human MB cells in the brain, consistent with the notion that only the innate, non-specific immune system (*e.g.*, microglia) is already active at this time point^22^. In contrast, the adaptive immune response (*e.g.*, lymphocytes) matures postnatally^23^. Interestingly, approximately 50% of injected brains still exhibited MB tumor foci up to P60 (Figure 1C), in contrast to our previous observations with human GBM, which typically regresses around P28 in the same model^17^. This finding aligns with studies demonstrating that SHH-MB tumors possess distinct immunosuppressive features^24^. Analyses of the xenotransplanted brains at E18.5, P2, and P7 revealed the presence of Tx in the cerebellum as well as in additional regions (Figure 1D, *red arrows*), suggesting that DAOY cells do not retain the anatomical-area identity in this model. Moreover, the size of Tx showed a marked increase over time (Figure 1D), and the engrafted cells were also positive for the proliferation marker Ki67, confirming their proliferative capacity at these time points (Figure 1E). The integration, persistence, and proliferation of xenotransplanted cells indicate that this model may be used to investigate the tumorigenesis and progression of MB.

### MB xenografts develop a tumor-associated vasculature, including a perivascular stem cell niche

MB development critically depends on the tumor microenvironment (TME), particularly on blood vascularization, which sustains the high metabolic demand of tumor cells and angiogenesis and correlates with tumor malignancy^25^. To ascertain whether human MB Tx develops a functional vascular system, we investigated the immuno-reactivity of Claudin-5, the most abundant tight junction protein of the Blood-brain Barrier (BBB)^26^, and Laminin, a key component of the basal lamina surrounding blood vessels^27^. Using an immunohistochemical approach, we detected Claudin-5 and Laminin within the Tx masses at both E18.5 and P7, exhibiting higher density and disorganization in the proximity of the Tx (Figure 2A). This observation aligns with previous findings, which demonstrated a significant disorganization of the vascular network within tumors compared to non-neoplastic tissues^27^. The co-localization of Laminin and Claudin-5 as early as E18.5 (Figure 2A) suggests the presence of a functional vasculature associated with the Tx. The presence of vessels co-stained for Laminin and Claudin-5 crossing tumor borders (Figure 2A, *higher magnification*) further supports this assumption, suggesting the establishment of vascular connections with the host tissue. To further characterize the TME, we assessed immunoreactivity for Vascular Endothelial Growth Factor (VEGF) and Cluster of Differentiation 31 (CD31), markers of angiogenesis and endothelial cells, respectively, which are also used in clinical settings to evaluate vascularity in solid tumors^25^. VEGF and CD31 were detected in the Tx at E18.5 and P7 (Figure 2 B, C). Remarkably, at P7, we found CSCs in the Tx as indicated by immunostaining for a human-specific anti-Nestin (hNestin) antibody^28^. Interestingly, hNestin^+^ cells colocalized with the vascular endothelial marker CD31 (Figure 2D), in line with a previous study on another *in vivo* model of human MB^29^. These results indicate that in the context of the embryonic brain, human MB Tx develops a functional vasculature and, likely, a perivascular CSC niche.

**Figure 2.**
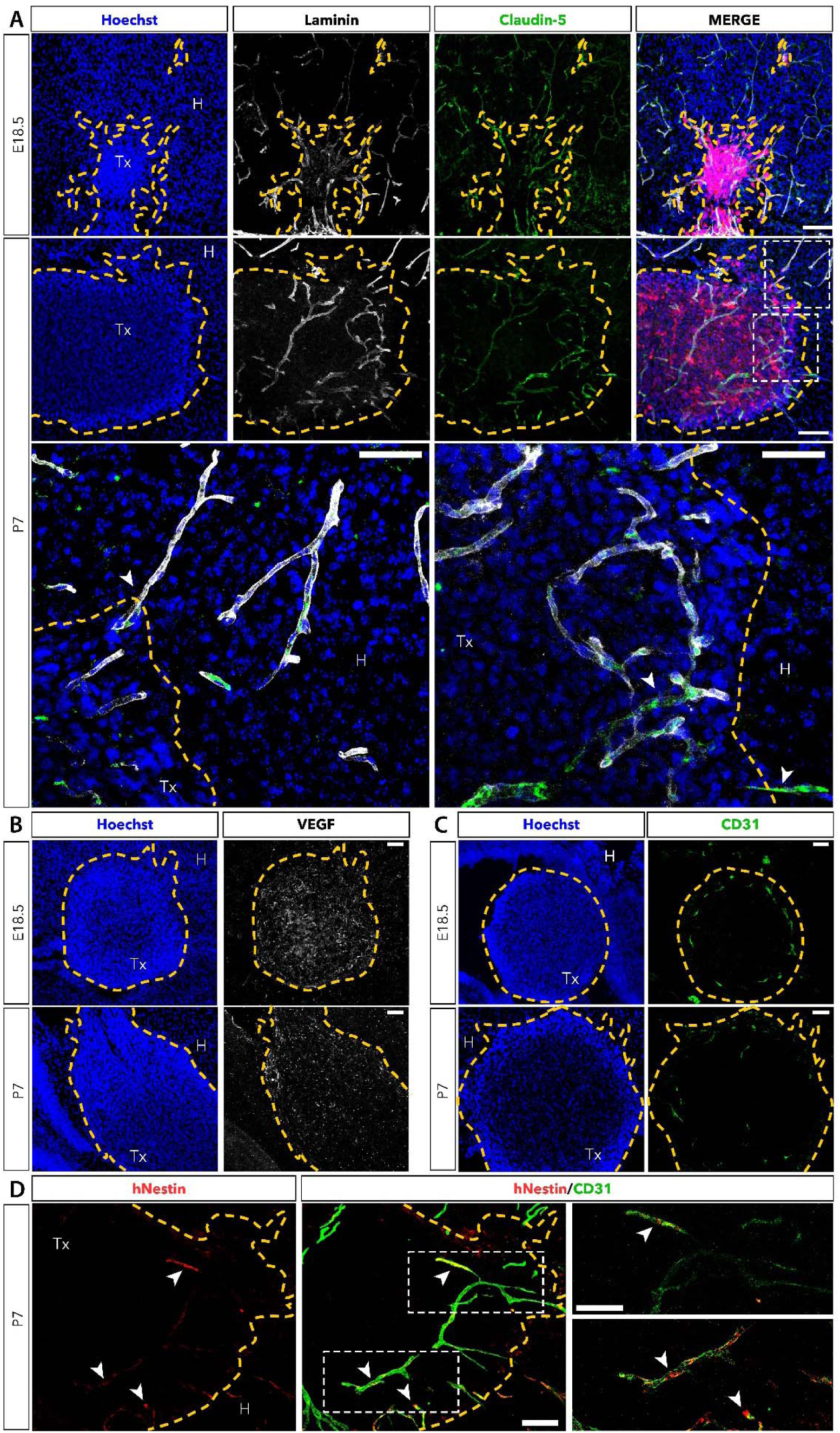
MB xenografts develop a tumor-associated vasculature including a perivascular stem cell niche. Immunofluorescence images of cryosections through xenotransplanted brains at the indicated time points. (A) Laminin and Claudin-5 signals colocalize at E18.5 and P7. Scale bar: 100µm. Boxes images show at higher magnification of double-positive Laminin/Claudin-5 vessels crossing the tumor borders (arrowheads) in the host tissue (left), or in the Tx core (right); scale bar: 100µm. (B) Immunofluorescence staining for VEGF inside the Tx. Scale bar: 100µm. (C) Immunofluorescence staining for CD31 inside the Tx. Scale bar: 100µm. (D) Immunofluorescence staining for human-specific Nestin (hNestin) and its colocalization with CD31 (arrowheads). Scale bar: 100µm. Dashed lines indicate Tx borders.

### MB xenografts trigger microglia infiltration and activation

Tumor-infiltrating cells, including microglia (MG) and tumor-associated macrophages (TAMs), significantly influence tumor progression by directly degrading the extracellular matrix or by suppressing T-cell activity, in turn, modulating angiogenesis and the establishment of an immunosuppressive TME^30^. These myeloid cells are particularly crucial in the SHH subgroup, as indicated by their increased infiltration compared to other MB subgroups^24^. However, the specific contribution of MG and TAMs is difficult to study in conventional immune-suppressed mice models, which fail to recapitulate the adaptative immunity biology^31^. Taking advantage of the WT background of our murine model, we analyzed the recruitment of MG/TAMs populations within the Tx masses. Immunostaining for the ionized calcium-binding adapter molecule 1 (IBA1) was evaluated as a marker of both MG and TAMs. At E18.5, we found IBA1^+^ cells only outside the Tx masses (not shown); however, at P7, we found IBA1^+^ cells also inside the Tx masses (Figure 3A, *arrows*). Interestingly, double staining for IBA1 and Cluster of Differentiation (CD68) (a marker for active MG and TAMs, when co-localized with IBA1^32^) showed the presence of active MG/TAMs inside the Tx masses (Figure 3A, *arrows*). These results indicate that the MB tumor is dynamically colonized by host myeloid cells.

**Figure 3.**
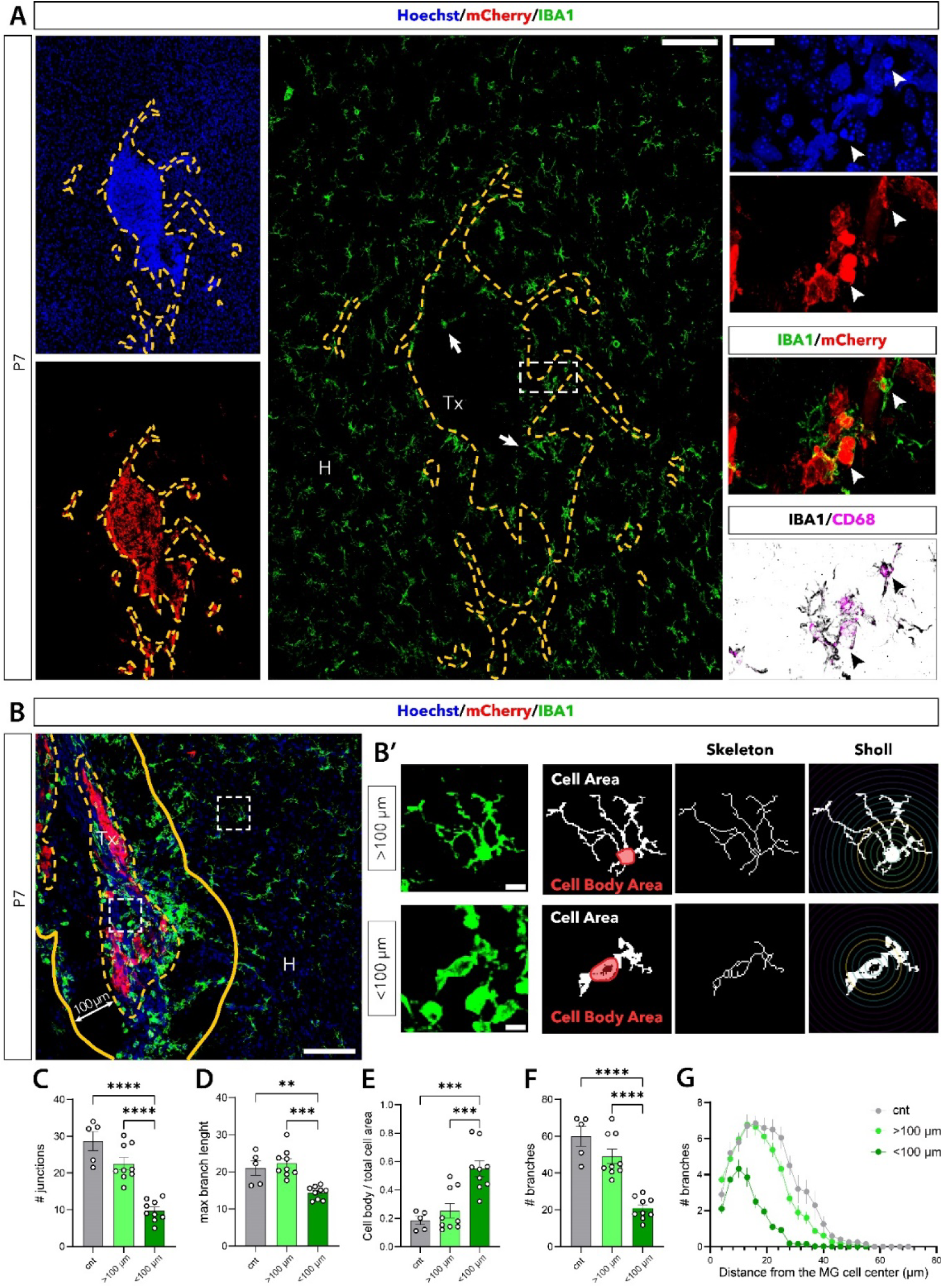
MB xenografts trigger microglia infiltration and activation. Immunofluorescence images of cryosections through xenotransplanted brains at the indicated time points. (A) Immunofluorescence staining for IBA1 showing MG/TAMs inside the Tx (arrows). Scale bar: 100µm. Co-localization of CD68, IBA1 staining with mCherry^+^ cells in the Tx region marked by the box and shown at higher magnification in the panels on the right. Arrowheads indicate CD68 and IBA1 double-positive cells colocalizing with disrupted mCherry^+^ nuclei. Scale bar: 20µm. (B) Immunofluorescence image showing IBA1^+^ MG/TAMs inside the Tx core (dashed line), or in the region surrounding the Tx (<100 µm, area between the dashed and plain lines) and >100µm from the Tx. Scale bar: 100µm. Boxes indicate the cells shown at higher magnification in the panels on the right. (B’) Maximum intensity projection (left) of IBA1^+^ cells in the indicated locations. Binarized and skeletonized schemes of the cells used for the quantification of the cell body/total cell area (E), number of junctions (C), maximum branch length (D), number of branches (F) and their distance from the cell center (G). Scale bar: 10µm. Data are represented as mean ± SEM (n = 4 cells from 4 animals per group; statistical significance revealed by using one-way ANOVA and Tukey’s multiple comparisons test; **p<0.01, ***p<0.001.

Furthermore, co-localization of IBA1^+^/CD68^+^ cells with mCherry^+^ nuclei (Figure 3A, *panels on the right*), suggested active phagocytosis of MB cells by the host myeloid cells. This observation was further corroborated by disrupted mCherry^+^ nuclei associated with IBA1 and CD68 double-positive cells (Figure 3A, *arrowheads*). To further validate the activation of MG/TAMs in association with MB foci, we analyzed the morphology of IBA1_ cells, distinguishing between their ramified morphology, indicative of a resting state, and an ameboid morphology, characteristic of activation^33^. In P7 brain slices, IBA1_ cells located within the Tx and within a 100 μm radius surrounding it were classified as MB-associated MG/TAMs. Their morphology was compared to that of IBA1_ cells situated more than 100 μm away from the Tx in the same section and to cells in brains from littermates not subjected to xenotransplantation (controls) (Figure 3B). We quantified specific morphological parameters, including the number of junctions (*i.e.*, points where cell processes branch), the number of branches (*i.e.*, total individual processes), and the maximum branch length (*i.e.*, the length of the longest process of a single cell). These analyses revealed that IBA1_ MG/TAMs associated with MB tumors exhibited an ameboid, activated morphology—evidenced by a significant reduction in the number of junctions, branches, and maximum branch length, and by increased cell body/total cell area ratio, compared to MG cells located outside the tumor and in control samples (Figure 3C-F). Finally, Sholl analysis—a widely used method to quantify cell branching complexity^34^–demonstrated a marked reduction in process complexity in Tx-associated MG/TAMs cells compared to that of cells outside the Tx and in controls (Figure 3G). These findings indicate that host myeloid cells are recruited and activated by the tumor grafts, supporting the utility of this model for studying tumor-immune interactions.

### Restoration of underexpressed miRNAs in MB suppresses tumor growth both *in vitro* and *in vivo*

Deregulation of miRNA expression is a hallmark of MB. We focused on a specific Pool of miRNAs (hsa-miR-124-3p, hsa-miR-127-3p, hsa-miR-134-5p, hsa-miR-135a-5p, hsa-miR-139-5p, hsa-miR-218-5p, hsa-miR-370-3p, hsa-miR-376b-3p, hsa-miR-382-5p, hsa-miR-411-5p, and hsa-miR-708-5p), which synergistically promote neural differentiation of non-neoplastic NPCs^35^, and were previously demonstrated to exert a tumor-suppressive effect in GBM^36^, another brain cancer characterized by CSCs^37^. MiRNA levels were investigated in two well-characterized human MB cell lines—DAOY and D283— representative of the SHH and Group 3/4 subgroups of this cancer. These cell lines exhibit distinctive transcriptional landscapes (Supplementary Figure S1), consistent with the significant molecular heterogeneity observed within these MB subgroups^7^. Small RNA-seq profiles indicated that about 30% of the miRNAs were deregulated in these MB cells, compared to their expression in the non-neoplastic cerebellum (GEO #GSE181520; Figure 4A). Next, we focused on the expression of the Pool in MB cells. Since this Pool was originally characterized in mouse NSCs^35^, the three human homologs of mmu-miR-376b-3p (the only miRNA of the Pool not conserved in humans) sharing similar seed regions were considered in the analysis (*i.e.,* hsa-miR-367a-3p; hsa-miR-367b-3p and hsa-miR-367c-3p, Supplementary Table S1). This analysis revealed that all eleven miRNAs of the Pool were underexpressed in cancer cells from both MB subgroups (Figure 4A).

**Figure 4.**
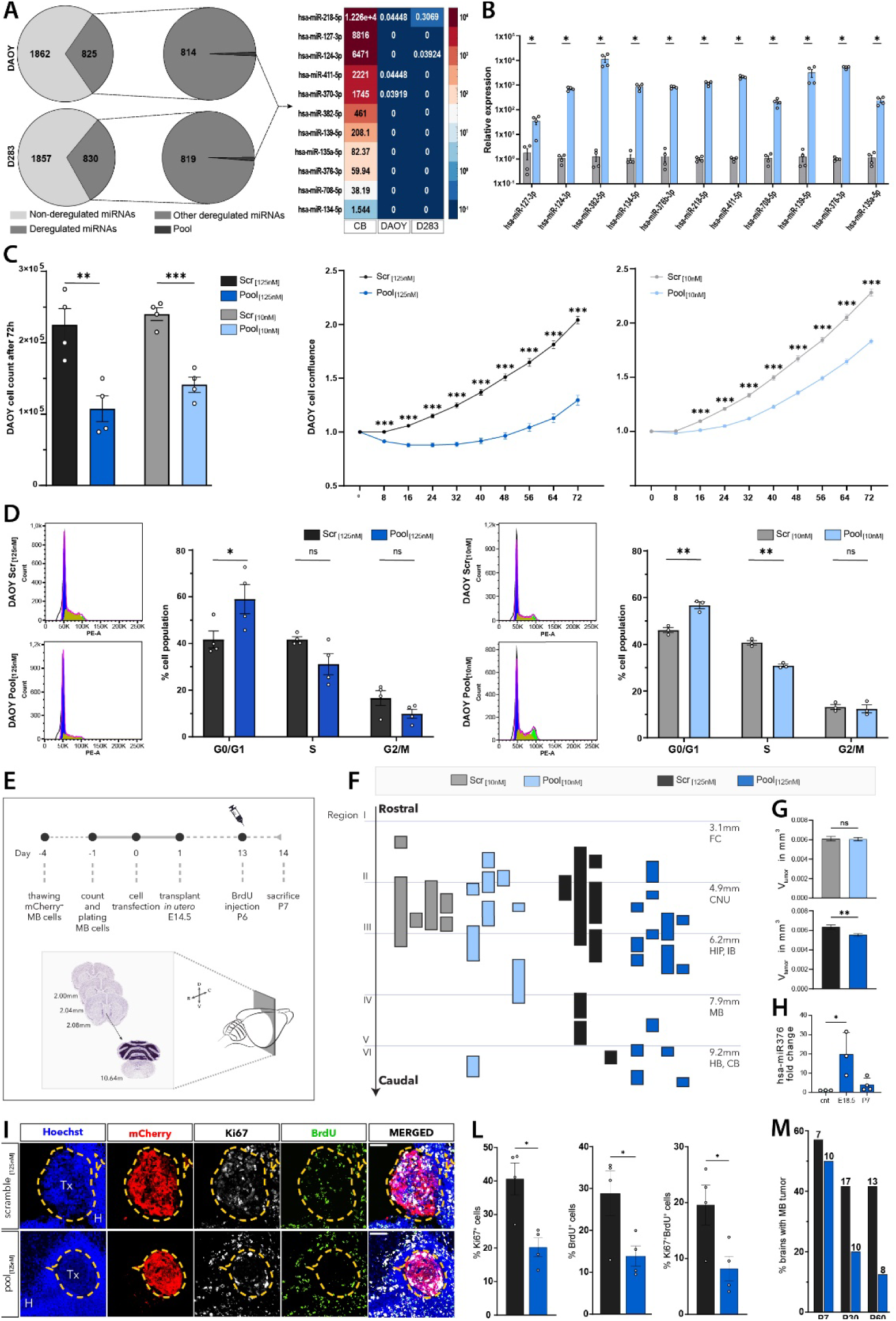
Restoration of underexpressed miRNAs in MB suppresses tumor growth both *in vitro* and *in vivo*. (A) Pie Plots showing the number of detected and deregulated miRNAs in small RNA-seq from DAOY and D283 MB cells and heatmap of the eleven deregulated miRNAs compared to their expression in human non-neoplastic cerebellum (CB); data are shown as transcripts per million (TPM). (B) QPCR mediated quantification of the eleven miRNAs upon transfection in DAOY cells, compared to their levels in Scrambled-transfected cells (control), at 10nM. Data are represented as mean ± SEM on n=4 biological replicates through a multiple t-test. (C) Quantification of cell proliferation upon the transfection of the Pool or Scrambled RNA at the indicated concentrations. Left: bar plot indicating the cell count seventy-two h after transfection. Centre and right: live imaging-mediated quantification of cell confluence over time. Statistical analyses were performed through a t-test for the cell count (n=4 biological replicates) and two-one-ANOVA for the cell confluence (n=6 biological replicates). (D) Representative cell cycle analysis of propidium iodide staining by flow cytometry seventy-two h after the transfection of DAOY cells with the Pool or Scrambled RNA at the indicated concentrations. Bar plots indicate the percentage of cells in the three cell cycle phases. Data are represented as mean ± SEM of n=4 biological replicates. Multiple t-test. (E) Scheme of the experimental procedure: DAOY cells, pre-transfected *in vitro* with the indicated concentrations of Pool or Scrambled RNA, were injected *in utero* at E14.5; BrdU was injected the day before the sacrifice (P6) and serial brain sections were collected. (F) Scheme showing the reconstructed Tx masses across serial section of the xenotransplanted brains (each column represents Tx masses from one brain), originated from mCherry^+^ DAOY cells transfected with the Pool or Scrambled RNA at the indicated concentrations. (G) Quantification of the Tx volumes. Data are represented as mean ± SEM of n = 4 biological replicates; t-test. (H) QPCR mediated quantification of hsa-miR-376b-3p in Tx masses isolated from transplanted brains. Data are represented as mean ± SEM of n=3 biological replicates at the indicated time points. One-way ANOVA. (I) Immunofluorescence staining for Ki67 and BrdU in Tx, originated from Pool- or Scrambled-transfected mCherry^+^ DAOY cells, in brains of P7 mice. Scale bar: 100 µm. (L) Percentage of Ki67^+^, BrdU^+^ and Ki67/BrdU double-positive DAOY cells in each Tx. (M) Percentage of brains with Tx over the total injected brains, originated from Pool- or Scrambled-transfected mCherry^+^ DAOY; sample size is reported at the top of each histogram bar. Data are represented as mean ± SEM of n=4 biological replicates. t-test test. *p<0.05, **p<0.01, ***p<0.001.

To investigate the biological effect of the Pool in MB, we transfected DAOY and D283 cells with two concentrations (10 nM or 125 nM) of the Pool—comprising eleven synthetic mimics in equimolar amounts—or with the same concentrations of a Scrambled RNA (control). Seventy-two h later, qPCR confirmed increased levels of all eleven miRNAs in DAOY (Figure 4B) and D283 cells (Supplementary Fig. S2), compared to Scrambled RNA-transfected cells. By unbiased cell counting, we found that the Pool significantly reduced the DAOY (Figure 4C, *left panel*) and D283 (Supplementary Figure S2) cells, compared with controls. Similarly, both Pool concentrations slowed DAOY cell growth over seventy-two h compared to the control, as indicated by a lower rate of cell confluence documented by label-free automated live imaging (Figure 4C, *right panel*). To confirm the antiproliferative effect of the Pool, seventy-two h after transfection with both concentrations, we performed Propidium Iodide (PI) staining and flow cytometry analysis in DAOY and D283 cells. We found a slower cell cycle in Pool-transfected DAOY (Figure 4D) and D283 (Supplementary Figure S2) cells at both concentrations, with a higher proportion of cells found in the G0/G1 phase and a lower proportion of cells in the S phase after the transfection of the Pool, compared to controls. Importantly, no apoptosis was observed upon transfection of both Pool concentrations in DAOY cells (Supplementary Figure S3), indicating that restoring these miRNAs was not toxic for these cells.

To investigate whether all eleven miRNAs are required to elicit the anti-proliferative effect of the Pool, we performed miRNA deconvolution experiments in DAOY cells, which are more responsive to the Pool than D283 cells. As expected, the complete Pool significantly slowed the growth of DAOY cells over seventy-two h compared to Scrambled RNA-transfected cells. In contrast, only hsa-miR-124-3p, when transfected alone, could phenocopy the effect of the complete Pool (Supplementary Figure S4A). However, seven of the eleven sub-pools – each lacking a different single miRNA – lost the effect of the complete Pool (Supplementary Figure S4B). These findings suggest that, while some miRNAs may contribute more significantly than others, the full anti-proliferative response likely relies on a synergistic interaction among the eleven miRNAs, consistent with previous reports on the cooperativity of the same pool of miRNAs in non-neoplastic NSC^35^, and in GBM^36^. Further studies will be required to address whether a potential hierarchy among individual miRNAs within the Pool can be leveraged for personalized MB treatments.

To investigate the tumor-suppressive effects of the Pool *in vivo*, mCherry_ DAOY cells, transfected with either the Pool or Scrambled RNA (at 10 nM or 125 nM), were injected into the embryonic telencephalon of E14.5 WT mouse embryos via *in utero* transplantation and injected mice were sacrificed at P7. The Tx masses were reconstructed with confocal images across serial sections in each mouse, and their volumes were quantified (Figure 4E). Transfection of the Pool at 125 nM in mCherry^+^ DAOY cells reduced the Tx mass volume, whereas 10 nM had no significant effect compared to Scrambled-transfected Tx (Figure 4F, G). Henceforth, the higher concentration of the Pool was selected for subsequent *in vivo* experiments. To quantify the stability of the miRNA mimics *in vivo*, mCherry^+^ Tx masses were collected by manual dissection from the slices at E18.5 or P7 (*i.e.,* 4 days and 2 weeks after cell injection, respectively). The abundance of the hsa-miR-376b-3p (*i.e.,* the only human-specific miRNA of the Pool) was quantified by human-specific primers for qPCR^36^. At E18.5, the hsa-miR-376b-3p level was significantly increased in Pool-transfected Tx masses compared to Scrambled-transfected Tx. However, the level of this miRNA returned almost to the basal at P7 (Figure 4H). Assuming similar stability for all the transfected mimics, this result indicates that the miRNAs are likely depleted within two weeks *in vivo*.

To validate the antiproliferative effect of the Pool *in vivo*, we quantified cells immunostained for the proliferative markers Ki67 and BrdU (administered twenty-four hours before sacrifice; Figure 4E) in tumors generated by Pool- or Scrambled-transfected mCherry_ DAOY cells. The Pool significantly reduced tumor cell proliferation *in vivo*, as indicated by a lower percentage of Ki67^+^, BrdU^+^, and Ki67^+^BrdU^+^ double-positive cells in the Tx masses (Figure 4I, L). Interestingly, the proportion of brains showing Tx was similar at P7 upon engraftment of both Pool- or Scramble-transfected mCherry^+^ DAOY cells, but this proportion dropped at both P30 and P60 in Pool-compared to Scrambled-transfected Tx (Figure 4M). Together, these results suggest that restoring the Pool suppresses MB growth and likely inhibits its progression in vivo but not its initiation.

### Restoration of the Pool suppresses MB migration and invasiveness

The poor prognosis of MB patients is largely due to its high invasive potential that, when leading to metastases, significantly increases the risk of tumor recurrence^38^. To assess the efficacy of the Pool in this process, we transfected DAOY cells with the lowest effective Pool concentrations and investigated migration *in vitro* and invasiveness *in vivo*. A scratch-wound assay in a label-free automated live imaging setup was used to monitor cell colonization of the wound for seventy-two h. The Pool significantly impaired the migration of MB cells, slowing the wound colonization with the earliest effects detectable starting from 32h (Figure 5A). Since the doubling time of DAOY in our culture conditions is 39.56 h, this result suggested that the Pool is likely anti-migratory *in vitro*. Transcriptomics analysis of these cells corroborated this result by showing the downregulation of genes belonging to the GO-term “positive regulation of cell migration pathway” (*i.e*., GO: 0030335) in Pool-transfected compared to Scrambled-transfected DAOY (Figure 5B). A similar suppression of migration-related genes was observed in Pool-transfected D283 cells (Supplementary Figure S5), though the suppressed genes differed from those in DAOY.

**Figure 5.**
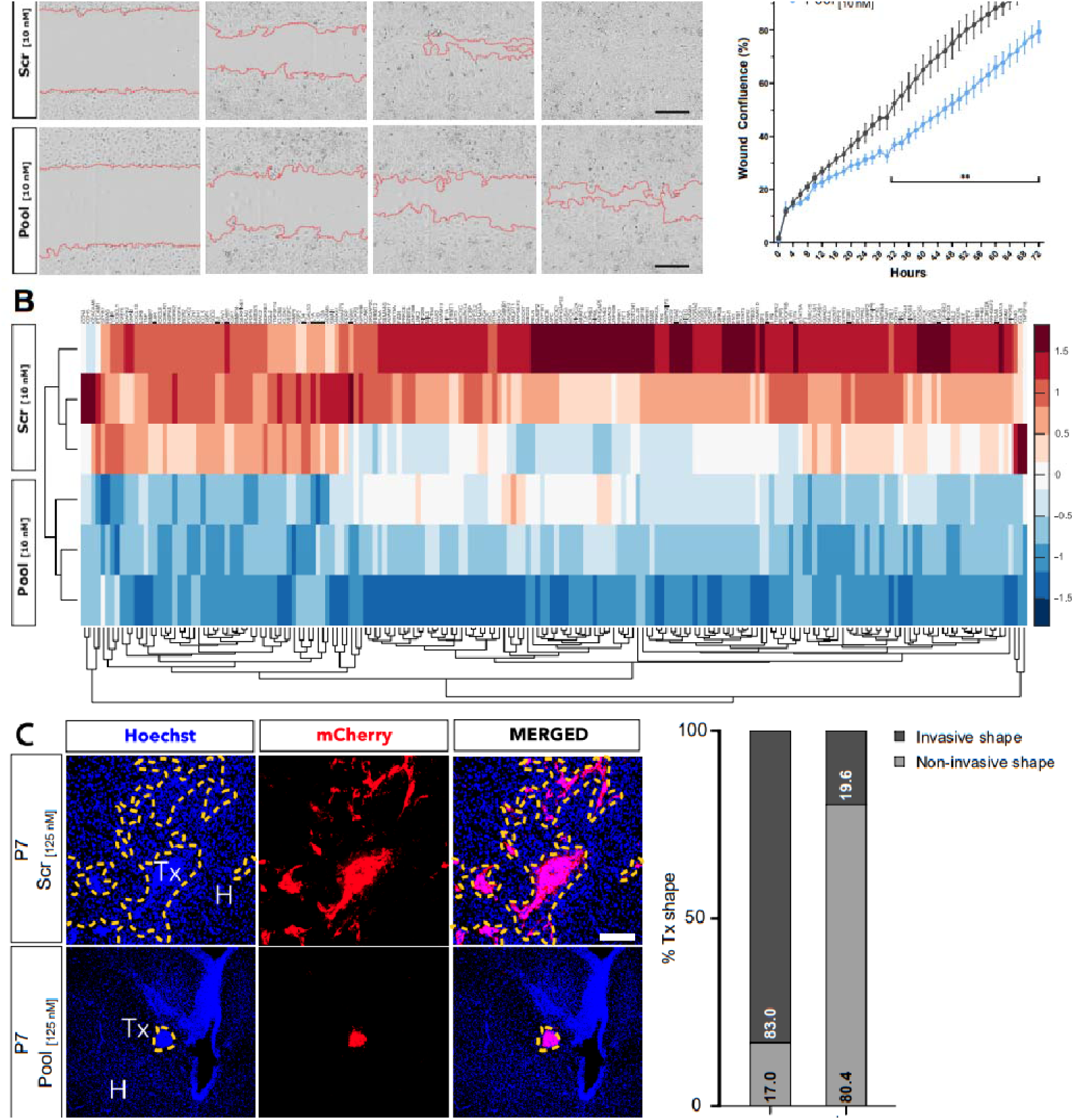
Restoration of the Pool suppresses MB migration and invasiveness. (A) Representative images of scratch-wound healing assay and quantification of cell migration of Pool- or Scrambled-transfected DAOY cells at the indicated concentrations, acquired by automated live imaging at the indicated time points. Scar bar: 300 µm. Data are represented as mean ± SEM of n= 8 biological replicates. One-way ANOVA. (B) Heatmap of poly(A) RNA-seq, showing levels of genes encoding for proteins involved in positive regulation of cell migration pathway (GO: 0030335) in DAOY cells, seventy-two h after transfection with Pool or Scrambled RNA. Samples were clustered based on biological samples (rows) and gene expression (columns). Only significantly downregulated genes are plotted. Wald test (n= 3 biological replicates). (C) Immunofluorescence images of cryosections through xenotransplanted brains at P7 showing representative examples of Tx with “invasive” vs non-invasive morphology of Tx originated from Pool- or Scrambled-transfected mCherry^+^ DAOY cells. Quantification is represented as a percentage of the two morphological categories relative to the total number of Tx (n=46 Pool; n=53 Scrambled). Scale bar: 100 µm. *p<0.05, **p<0.01, ***p<0.001.

To investigate the efficacy of the Pool on MB invasiveness, we analyzed the morphology of Tx masses at P7 *in vivo*. Morphological assessment revealed that 80% of the tumor masses generated from Pool-transfected mCherry_ DAOY cells exhibited a non-invasive phenotype characterized by expansive, bulky growth patterns with a well-defined, localized center. In contrast, 83% of the tumors derived from Scrambled-transfected cells displayed a distinct invasive phenotype, marked by infiltrative growth patterns with irregular borders (Figure 5C). Collectively, these results demonstrated that the Pool exerts multimodal tumor-suppressive effects, effectively limiting MB proliferation and invasive potential.

### The eleven miRNAs of the Pool converge on the same genes in different MB subgroups

To infer the possible mechanism underlying the tumor-suppressive effects of the Pool in MB, we performed poly(A)-seq and Liquid Chromatography/Mass Spectrometry (LC/MS) proteomics analyses (Figure 6). Pool- and Scrambled-transfected DAOY and D283 cells were harvested, and RNA was extracted. Principal component analysis (PCA) of transcriptomics data indicated a distinct segregation of Pool- vs Scrambled-transfected samples along the PC1 and PC2 in both cell types (Supplementary Figure S6). Transcriptomic analyses identified 6,712 and 6,542 genes significantly altered by the Pool in DAOY and D283 cells, respectively (Supplementary Tables S5-S6), suggesting a comparable transcriptional response in both MB groups. To identify the mechanisms underlying the effects of the Pool in MB, we focused on the downregulated transcripts since miRNAs primarily act as repressors. GO analysis of significantly downregulated transcripts revealed an enrichment of genes belonging to pathways related to proliferation, cell cycle, and migration in both DAOY and D283 cells (Figure 6A-B, *arrows*), consistent with the observed effects of the Pool. Remarkably, 771 transcripts were downregulated in both DAOY and D283 cells. Many of these genes belonged to cell cycle-associated GO categories. They linked to MB pathogenesis—reported as genetic variants, differentially expressed genes, or established oncogenic drivers in pediatric and adult MB (Figure 6C). These results suggested that the Pool might exert similar functions in different MB groups by converging on a shared set of genes. Since miRNAs primarily exert their repressive functions at the post-transcriptional level^39^, we corroborated this finding with proteomic analyses. Principal component analysis (PCA) of the proteomics analysis confirmed the distinct separation between the Pool- vs. Scrambled-transfected cells along the PC1 and PC2 (Supplementary Figure S7). The Pool significantly altered 508 (8.14%) out of the 6235 proteins identified in DAOY cells (Supplementary Table S7). Interestingly, 158 of the 230 downregulated proteins were also found to be downregulated at the transcriptional level (Figure 6D). Thus, we prioritized these putative targets in the subsequent analyses. As expected, protein-protein interaction (PPI) analysis revealed that these 158 proteins are functionally interconnected, with a significant enrichment (p-value < 10e-16). Functional clustering of the downregulated proteins via the STRING algorithm grouped the downregulated proteins into five functional clusters related to “cell adhesion/migration”, “RNA splicing/metabolism”, “catabolic/biosynthetic processes”, “transcription”, and “chloride channel complex” (Figure 6D). These results are consistent with the anti-migratory and anti-proliferative role of the Pool in MB.

**Figure 6.**
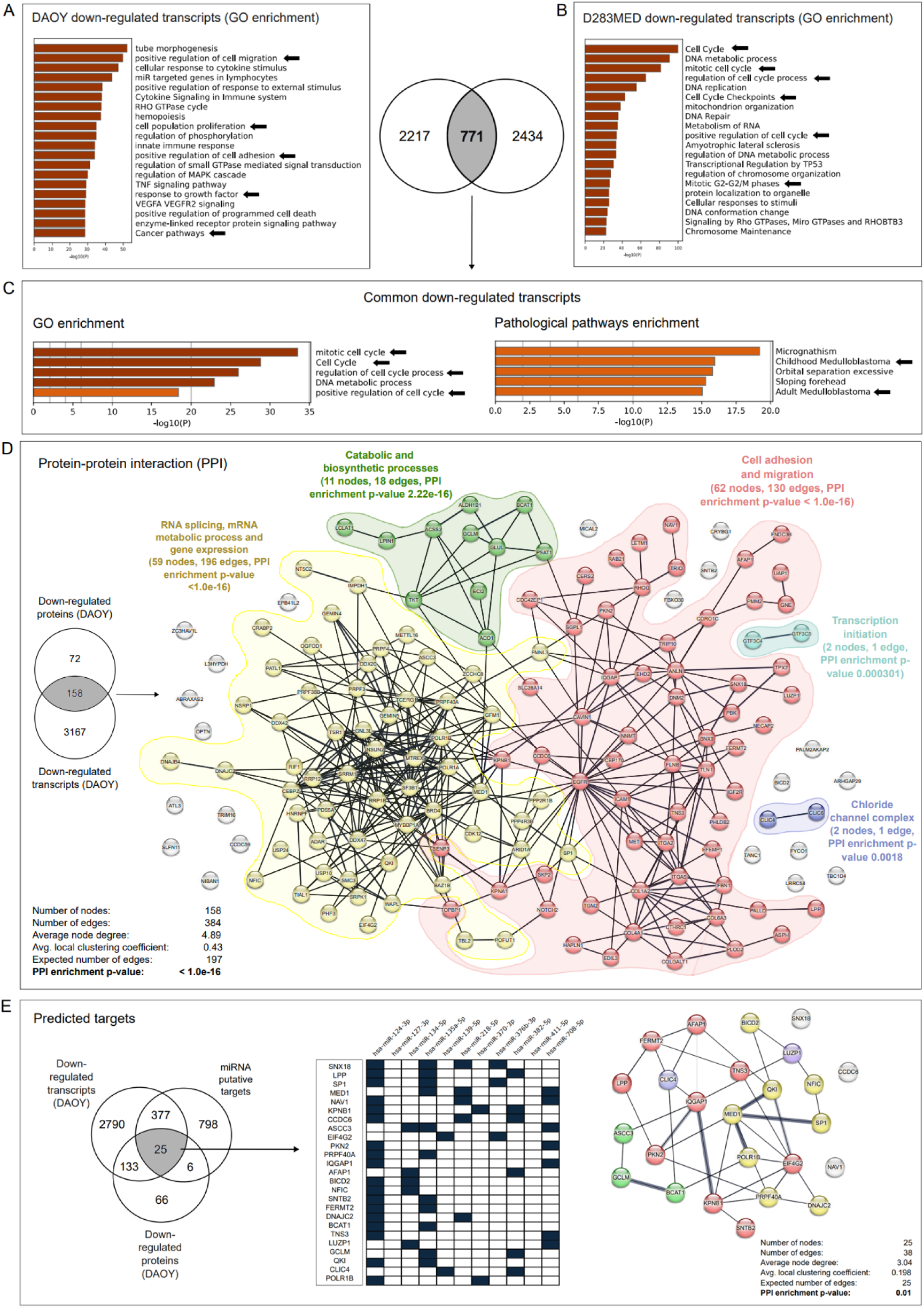
The eleven miRNAs of the Pool converge on the same genes in different MB subgroups. (A, B) GO analysis of the top non-redundant enrichment clusters of the downregulated genes in the indicated MB cells and Venn diagram showing number of common and unique genes. (C) GO analysis of the top non-redundant enrichment clusters (*left*) and gene-disease association network (*right*) of the common downregulated transcripts. Color scale reflects the statistical significance of the enrichment. (D) Protein-protein interaction (PPI) analysis of the down-regulated targets at both transcript and protein levels in Pool-transfected DAOY cells. Colors reflect the biological functions of the protein clusters. Statistical significance was determined by STRING PPI analysis. (E) Venn diagram showing the intersection of the predicted miRNA targets with the downregulated genes upon Pool transfection (*left*), heatmap showing miRNAs that are predicted to interact with the targets (*center*) and PPI analysis of the shared targets in DAOY cells. The color code is related to the PPI in C. Statistical significance of DE transcripts/proteins on n=3 (transcriptomics) and n=5 (proteomics) biological replicates. Parametric t-test. *p<0.05, **p<0.01, ***p<0.001.

To identify which of the downregulated proteins were possibly subject to miRNA-mediated repression, we employed the DIANA-microT-CDS algorithm^38^ for all the putative targets of each of the eleven miRNAs of the Pool (Figure 6E). Given the functional redundancy among these miRNAs, we focused on targets shared by at least two of the eleven miRNAs and cross-referenced these with genes showing downregulation at both transcript and protein levels, upon the transfection of the Pool. This analysis identified 25 putative targets (*i.e.,* SNX18; LPP; SP1; MED1; NAV1; KPNB1; CCDC6; ASCC3; EIF4G2; PKN2; PRPF40A; IQGAP1; AFAP1; BICD2; NFIC; SNTB2; FERMT2; DNAJC2; BCAT1; TNS3; LUZP1; GCLM; QKI; CLIC4; POLR1B) (Figure 6E). Remarkably, all these targets were significantly interconnected according to PPI analysis (enrichment p-value = 0.01) and emphasized a set of proteins known for their involvement in “cell migration and invasion” (Figure 6E, *red*). As repression of these putative targets mediates the tumor-suppressive effects of the Pool in different MB subgroups, we propose these proteins likely represent promising candidates for future therapeutic development.

## DISCUSSION

In this study, we present a miRNA-based therapy targeting human MB and validate its efficacy in a novel embryonic preclinical model that mirrors the tumor’s development and progression. We demonstrate that the Pool significantly reduces MB growth and invasiveness *in vivo*. Finally, through multi-omics analyses, we identify the genes and pathways underlying the tumor-suppressive effects of the Pool, revealing potential therapeutic targets for further exploration in precision medicine.

The initial characterization of our preclinical model indicates that human MB cells can initiate tumors in the brain of WT mouse embryos. This model allows to recapitulate key aspects of MB biology within a developmental time window that closely mirrors the pathophysiology of this pediatric cancer, overcoming key challenges in preclinical MB research. In fact, traditional murine MB models with syngeneic mutations have been valuable for testing hypotheses related to tumor initiation and progression. However, modelling murine MB neither accurately replicate the human-specific cells/signaling pathways involved in MB initiation, nor the neuro-immune interactions that mediate its progression in patients. A widely used alternative, orthotopic xenotransplantation of human MB cells into adult animals, also has significant limitations. Firstly, the adult nervous system lacks the widespread expression and roles of the embryonic signaling pathways (such as SHH and WNT), which are critical for MB (re)formation. Secondly, adult animals that are deliberately immune-compromised to prevent xenograft rejection, fail to mimic the neuro-immune interactions that influence drug toxicity and efficacy. Hence, these models are unsuitable for testing immune-targeted therapies, such as CAR-T treatments, or cancer vaccines. Unsurprisingly, most oncology drugs fail in late-stage clinical trials due to a lack of efficacy in patients ^40^. Thus, we anticipate that our strategy of MB xenotransplant in WT mouse embryos may serve as a cost-effective and translationally relevant alternative to complement the existing models for preclinical studies. Through histological analysis of our preclinical model, here we demonstrate that Tx masses develop a TME that includes blood vessels, a vascular-associated cancer stem cell niche, and infiltrating MG/TAMs. While these findings confirm that our model enables the study of the cellular mechanisms underlying MB vascularization and myeloid cell infiltration *in vivo*, several key questions remain unanswered. For instance, it is unclear whether the tumor-associated vasculature in our model arises from co-option of host blood vessels or *de novo* angiogenesis from transplanted MB cells. Additionally, we have yet to determine whether the tumor-associated vessels develop a functional BBB. Addressing these questions will be essential for further validation of our model for preclinical studies, particularly in contexts where vascular permeability plays a critical role in drug delivery and therapeutic efficacy. Interestingly, the presence of human Nestin_ stem cells associated with tumor vasculature further highlights the potential of our model for developing therapies that target the CSC niche, as well as for identifying biomarkers secreted by these cells. Thus, our model could meaningfully contribute to theragnostic advancements targeting the CSC subpopulation—a critical driver of MB initiation and recurrence.

Finally, we provide the demonstration of a possible tumor suppressive gene therapy based on a specific Pool of miRNAs, *in vivo*. Strikingly, the restoration of the Pool suppresses MB growth and invasiveness, thus addressing major challenges associated with poor prognosis in MB patients. Interestingly, the eleven miRNAs comprising the Pool are underexpressed across multiple MB subtypes, suggesting their potential role as epigenetic drivers of tumorigenesis and/or progression. While numerous studies have established that altered miRNAs expression is a hallmark of MB, the combination of miRNAs subject of our study was not tested as a therapeutic strategy against this malignancy^41–45^. Moreover, most — if not all — of these studies have addressed miRNAs singularly, an approach that has led to clinical trial failures in other cancers^15^. Hence, this study is the first to demonstrate the efficacy of a miRNA combination against human MB *in vivo*. This result, together with our recent finding that the same Pool is tumor-suppressive in GBM^36^, raises the tantalizing possibility that these miRNAs could serve as an anticancer therapy effective across multiple brain malignancies characterized by a CSC niche.

Our results indicate that the rescued miRNAs remain stable *in vivo* for about two weeks. From a translational perspective, this stability aligns with the dosing schedule of other RNA-based drugs, such as Onpattro, which is administered every three weeks (https://www.onpattrohcp.com/dosing-administration). Although here we did not explore the direct administration of the miRNA Pool *in vivo,* the local injection of synthetic small RNAs is feasible in clinical settings, and offers several advantages, including potentially faster approval processes compared to viral or cell-based delivery systems. Future studies should explore local delivery methods, such as biodegradable implants^46^, and/or chemical modifications of the miRNA mimics, to prolong their therapeutic effects^47^.

Finally, here we infer the mechanisms underlying the tumor suppressive efficacy of the Pool in MB. Multi-omics analysis following restoration of the Pool revealed a set of genes modulated by the miRNAs in two of the most frequent and aggressive MB subgroups. Interestingly, these genes are highly interconnected and encode for proteins involved in growth regulation, cell division and cancer progression in many cancers, including MB (Figure 6D). In agreement, inhibition of the transcription factor SP1 by Tolfenamic acid reduces MB growth^48^, and inhibition of the phosphorylation of mediator complex subunit 1 protein (MED1) by antagonists of the protein kinase CK2 underlies the anticancer efficacy of this class of drugs in MB^49^. Moreover, the functional inhibition of the nuclear import receptor karyopherin β1 (KPNB1, a protein involved in the nuclear import of most proteins as well as in mitosis) induces growth arrest and apoptosis in GBM^50^. Finally, SP1 as well as the other targets of the Pool such as EGFR, CDK12, and BRD4 identified in our study have already been investigated as therapeutic targets in other cancers, both at clinical and pre-clinical levels^51–54^. Thus, the proteins mediating the tumor-suppressive effects of the miRNAs identified in our study represent compelling candidates for future investigation, with the potential to inform MB precision medicine and combinatorial therapeutic strategies.

## MATERIAL AND METHODS (detailed methods are available in the supplementary material)

### Cell cultures, and transfection

DAOY cell line or D283 cell line were maintained in DMEM (Sigma Aldrich) with 1% of penicillin-streptomycin (P/S) antibiotics (ThermoFisher), or EMEM (Sigma-Aldrich) respectively, supplemented with 10% Fetal Bovine Serum (FBS, Sigma Aldrich) heat-inactivated (56°C for 30 min) and 1% of L-glutamine (Lonza Srl). <u>Transfection</u>: Cells were transfected with either 10 nM or 125 nM of a miRNA Pool or Scrambled siRNA (miRIDIAN, Dharmacon) using Lipofectamine 2000 (Life Technologies) in RPMI-1640 MegaCell medium (Sigma-Aldrich) for 6 hours (h). For subPools containing 10 miRNAs, any missing mimic was replaced with Scrambled RNA to maintain equal final concentrations. Following transfection, cells were cultured in serum-free complete Neurobasal (NB) medium (Gibco) supplemented with 1% penicillin-streptomycin, 1% L-glutamine, and 2% B-27 (Sigma-Aldrich).

### Animals and surgery

Mice were housed under standard conditions at the IIT animal facility in Genova, Italy. All experiments and procedures were approved by the Italian Ministry of Health and IIT Animal Use Committee, according to the Guide for the Care and Use of Laboratory Animals of the European Community Council Directives. WT CD1 females and males were crossed, and mouse embryos were used for *in utero* xenotransplantation experiments at 14.5 days *post coitum* (dpc). For all time-mated animals, the vaginal plug day was defined as 0.5 (E0.5). <u>Surgery</u>: transfected mCherry^+^ DAOY cells were injected 24h after, at a final concentration of 50.000 cells/μL and 10% sterile Fast Green (Sigma Aldrich). The surgical procedure was performed as previously published^17^. <u>BrdU labeling</u>: Postnatal day 6 mice received a single intraperitoneal injection of bromodeoxyuridine (BrdU, 50 mg/kg, Sigma-Aldrich) dissolved in PBS. Mice were sacrificed by cervical dislocation at the indicated time points.

## ACKNOWLEDGMENTS

The authors wish to thank A. Simi for advice on the *in utero* surgery procedure; M. Morini, E. Petrini and D. Vozzi and the technical staff at IIT’s Animal, Neurofacility and Genomics facility for excellent support; D. Russo and I. Penna for the support in acquiring qPCR data. We also wish to thank all the colleagues at “the Neurobiology of miRNA lab” for advice and discussion. We apologize to those colleagues whose work could not be cited due to space limitations.

## AUTHOR CONTRIBUTION

Designed research and experiments (CS, LLR, RCP, DDPT). Performed experiments (CS, LLR, RCP, LG, RP, ADF). Data acquisition and analyses (CS, LLR, RCP, LG, ADF, AA). Drafted the manuscript (CS, LLR, DDPT). Supervision (RCP, GB, AA, KT, DDPT). Acquired funding (RCP, DDPT). All authors read, revised, and approved the final version of the manuscript.

## DATA AVAILABILITY

Transcriptomics and proteomics datasets generated in this study are available upon request

