## Supplementary Material and Methods and Figures S1-S7 and Tables S1-S4 for "A microRNA-based Therapy against Human Medulloblastoma Validated in a Xenotransplant Model using Wild-Type Mouse Embryos"

**SUPPLEMENTAL INFORMATION**

**CONTENT:**

- **Detailed materials and methods**
- **References for detailed materials and methods**
- **Supplementary figures S1-S7**
- **Supplementary Tables S1-S7**

**DETAILED MATERIALS AND METHODS**

**Cell cultures and *in vitro* assays.** DAOY cell line (MB subgroup SHH; ATCC, HTB-186), or D283 cell line (MB subgroup 3/4; ATCC, HTB-185) were maintained in DMEM (Dulbecco’s Modified Eagle Medium; Sigma Aldrich) with 1% of penicillin-streptomycin (P/S) antibiotics (ThermoFisher), or Eagle’s Minimum Essential Medium (EMEM, Sigma-Aldrich) respectively, supplemented with 10% Fetal Bovine Serum (FBS, Sigma Aldrich) heat-inactivated (56°C for 30 min) and 1% of L-glutamine (Lonza Srl). Stable expression of mCherry (DAOY mCherry^+^) was obtained by transduction with a pLV-CMV-LoxP-mCherry-LoxP- lentivirus (Plasmid #65726, Addgene) (MOI=5). Positive clones were selected by fluorescence-activated cell sorting (FACS) for 99% positive cells and expanded for subsequent experiments. Transfection: cells seeded in 12-well plates at a concentration of 6x10^4^ cells/well were transfected with the Pool, subPool of 10 miRNAs, each miRNA singularly, or scrambled RNA mimics (miRIDIAN, Dharmacon^®^, USA; reported in Table S1) at equimolar concentration to a final concentration of 125 nM (~11.4 nM each miRNAs) or 10 nM (~0.9 nM each miRNAs), with Lipofectamine-2000 (Thermofisher) in RPMI-1640 MegaCell medium (Sigma-Aldrich) for 6 hours. Cells were then cultured in serum-free, complete Neurobasal (NB) (Gibco) supplemented with 1% P/S, 1% L-glutamine and 2% B-27 (Sigma-Aldrich). In these growth conditions, cell doubling time was evaluated using cell-by-cell analysis of live images with an Incucyte^®^ S3 (Sartorius, Italy) quantitative live-cell imaging (<https://www.doubling-time.com/compute.php>). Cell counting: MB cells were seeded at 6x10^4^ cells/well in 12-multiwell plates and after 24 hours transfection assays were performed as described above**.** After 72 hours, cells were detached with trypsin-EDTA (Gibco 25200-056) at 37°C for 2-3 min. Trypsin activity was inhibited with DMEM supplemented with 10% FBS. Cells were centrifugated at 1200 rpm for 5 minutes at RT and resuspended in DMEM. After, 10μL of the cell suspension were mixed with 10μL of trypan blue and cells were counted by the aid of the cell counter DeNovix CellDrop BF. Cell cycle analysis: DAOY and D283 cells were seeded at a concentration of 4x10^5^ cells/flask in T25 flasks and transfected as described above with the Pool or Scramble RNA at 10 nM for 72h hours. Cells were detached as above and fixed in ethanol 70% at - 20°C. Suspensions were kept for 40 minutes on ice, pelleted at 1500 rpm for 5 min at RT and resuspended in 1mL of cold PBS for 15 minutes. Then, cells were pelleted again at the same conditions and resuspended in 500mL of propidium iodate (PI) solution (0.2mg/mL RNase-A, 0.05% Triton x-100 and 60mg/mL PI) for 1 hour at 37°C and resuspended in 400mL of PBS at RT. Flow cytometry was performed using the BD FACS ARIA III (BD Biosciences, San Josè, CA, USA) equipped with a forward scatter photomultiplier tube. Cells were scanned for PI with a yellow-green laser (561 nm) and detected by a 582/15 nm band pass filter, at a rate of 500 events/s. Forward scatter area (FSC-A) and side scatter area (SSC-A) were used to exclude debris and cell aggregates; doublets were excluded from the basal cell population by side scatter SSC-W vs SSC-H. Samples were analyzed with FlowJo_v10.8.0 (BD Biosciences) to study the cell cycle. Apoptosis assay: DAOY and D283 cells were seeded at a concentration of 4x10^5^ cells/flask in T25 flasks and then transfected as described above with the Pool or the Scramble RNA at 125 nM and 10 nM for 72 hours. Annexin V and PI staining was done with “Dead Cell Apoptosis Kit with Annexin V FITC & Propidium Iodide for Flow Cytometry” kit (Invitrogen, Cat V13242) following manufacturer's instructions. Samples were analyzed by measuring the fluorescence emission at 530 nm and > 575 nm, by FACS and analyzed as described above. Wound healing assay. DAOY cells were seeded the day before transfection at a concentration of 3x10^4^ cells/well in 96-multiwell plates (Incucyte^®^ image lock 96 well microplate), transfected with the Pool or with the Scramble RNA at 10 nM as described above and after 24 hours, 700-800μm wide wounds in the cell confluent monolayers were created using the 96-pin mechanical device - Wound Maker according to the manufacturer (Sartorius, Italy). Plates were analyzed using Incucyte^®^ S3 instrument using 10x objective with 2 images/well with a frequency of every 4 hours for 72 hours. The raw data were analyzed by calculating the average of the two acquisitions per well for each biological replicate, followed by normalization of the measurements to the 0-hour time point.

**Animals and surgery.** WT CD1 pregnant females were subject to the surgical procedure of *in utero* xenotransplantation experiments at 14.5 days *post coitum* (dpc, vaginal plug day was defined as 0.5). Surgery: mCherry^+^ DAOY cells were transfected with the Pool or Scramble RNA 24 hours before the transplantation. On the day of injection, cells were detached with trypsin (0.25%) and 50.000 cells/μL in PBS (without Mg^2+^ and Ca^2+^, Gibco) were injected as previously published^1^. BrdU labelling was done at the indicated time points via intraperitoneal (IP) injection of 50 mg/kg of bromodeoxyuridine (BrdU, Sigma-Aldrich) in PBS. Mice were sacrificed by cervical dislocation at the indicated time points.

**Immunofluorescence and analyses**. *In vitro.* Pool- or Scrambled-transfected mCherry^+^ DAOY cells were seeded on coverslip, fixed for 15 minutes with 4% PFA 72h after the transfection, permeabilized with 0,1% triton-PBS and antigen retrieval was performed for 10 minutes at 85°C. Cells were incubated in blocking solution for 1 hour at RT, incubated with the primary antibody (Table S3) O/N at 4°C for 24 hours. Cells were then incubated with the secondary antibody (Table S4) for 2 hours at RT. *In vivo:* brains with tumor engraftments (Tx, documented with an Olympus SZX16 or Olympus U-RFL-T stereoscope) were fixed overnight (O/N) at 4°C in 4% paraformaldehyde (PFA, Sigma-Aldrich) in PBS, de-hydrated in 30% sucrose solution and embedded in Optimal Cutting Temperature Compound (OCT). Coronal cryosections were cut at 20 μm, rehydrated in 1x PBS and (when specified) subjected to antigen retrieval with 10 mM citric acid at pH 6.0 for 10 min at 95°C and permeabilized with progressive steps in 0.3% and 0.1% Triton X-100 in 1x PBS (PBST). Slides were incubated in blocking solution (5% normal goat serum – NGS – in 0,1% triton-PBS) for 1h at room temperature (RT). Sections were incubated with the primary antibodies (Table S3) in blocking solution O/N at 4°C. For detection of BrdU incorporation, tissues were pre-treated with 2N HCl for 30 min at 30.2°C, washed 1x PBST at RT for 24 hours; slices were washed with 0.1% PBST and incubated with secondary antibodies (Table S4) in blocking solution for 2h at RT. Nuclei were counterstained with Hoechst 33258 (Sigma-Aldrich) at a final concentration of 5 µg/mL. Sections were mounted with ProLong Gold Antifade solution (Invitrogen). Microglia morphology analysis: images were acquired across 13 Z-stacks (1 μm in thickness) from Iba1-stained brain-positive slices. The acquisition was centered at the point of maximum intensity for mCherry signal. Imaging was performed using a Nikon A1 confocal laser-scanning microscope with a 20x objective. One randomly chosen slice per animal was acquired. For the calculation of the cell area and cell-body area, measurements were conducted using the freehand selection tool in ImageJ and measurement was performed using the ImageJ option. For Sholl analysis, cells were cropped and thresholded to produce a binary (black-and-white) image, which was manually refined for each cell and analyzed using the ImageJ plugin ShollAnalysis^2^. Starting radius was set at 4 μm, ending radius at 70 μm (radius step size: 3 μm). Parameters were set with a starting radius of 4 μm, an ending radius of 70 μm, and a step size of 3 μm. The same binary image was then used for skeleton analysis; skeleton was generated by the ImageJ Plugin Skeletonize^3^ and analyzed with the AnalyzeSkeleton plugin to quantify number and length of branches and number of junctions. Tx characterization: 40 μm coronal sections were obtained from at least 4 brains, progressing rostro-caudally while maintaining strict anatomical reference points. This ensured that each slice occupied the same position across all brains. Brain slices were categorized into six principal anatomical *regions* for ease of representation. A single brain slice possessing mCherry signal is considered a *tumor focus.* The set of consecutive brain slices presenting *tumor foci* (mCherry fluorescence signal) is defined as a *tumor mass* (or simply, *tumor*). Tumor volume: was measured in coronal sections of embryonic brains by the diameter (d) of each tumor in its greatest dimension and volume (V) was estimated with the following ellipsoid formula^4^: 𝑉 = 𝜋𝑎𝑏9, where *a* and *b* correspond to the sagittal and axial plane, respectively. MCherry fluorescence signal defined *tumor area.* Analysis was restricted to tumor masses that occupied at least 6 consecutive brain slices; n = tumor masses (data from a single brain were pooled considering treatment conditions and considered as a whole). Cell proportions measurements: for cell density within the tumor area, Hoechst^+^ nuclei were counted within tumor area (defined by mCherry fluorescence) and divided by the tumor area; proportion of cells were calculated as the number of cells Ki67^+^ or BrdU^+^ or Ki67^+^BrdU^+^ over total Hoechst^+^ nuclei within the tumor area and normalized to the tumor area. During the count, cells were considered also for their morphology to exclude mouse nuclei counting was performed by considering different regions of the tumor mass and different regions of the brain (when applicable); n = number of brains (average of values from single brains). Imaging. Immunofluorescence images on slides were acquired with Confocal Nikon A1; images were analyzed with Nikon software version 4.11.0 (NIS Elements Viewer) and processed with ImageJ version 1.48v (https://imagej.net/ij/, 1997-2018).

**RNA extraction, qPCR, sequencing.** RNA was extracted from cultured cells by QIAzol (Qiagen), from xenografts with QIazol and a TissueLyser II (Qiagen) with 5 mm stainless steel beads pre-treated for RNAse and DNAse, following 3 × 1 min cycles at 30 Hz of homogenization, according to the manufacturer’s instructions. Total RNA was treated with DNase I (Sigma) and then quantified using a Nanodrop ND-1000 spectrophotometer (Thermo Fisher Scientific). RNA purity was evaluated with the 260/280 and 260/230 absorbance ratios: a value of ~ 2.0 was considered indicative of good quality RNA. Quantification of the levels of the 11 miRNAs was performed with the miScript SYBR Green PCR kit (Qiagen) in a RTqPCR 7 Pro (02-st32.0); or by TaqMan^®^ Custom 384-well Cards (Applied Biosystem), in ViiA 7 Real-Time PCR System with TaqMan Array Block (Life Technologies). Expression levels of U6 snRNA were considered for relative quantification of miRNAs using the delta-delta Ct method. *In vivo* stability of hsa-miR-376b-3p was performed on RNA extracted from xenotransplanted MB Tx with qPCR and species-specific primers as previously reported in Rancati et al^5^. Poly(A)-seq: total RNA extracted from DAOY and D283 cells as described above was treated with DNAse I (Sigma, AMPD1-1KT). Quantity and quality of RNA were then measured by Qubit 4 Fluorometer (Thermo Fisher) and Bioanalyzer RNA chips (Agilent), RNA with RNA integrity number (RIN) values ≥ 7 were selected for the study. 100 ng of high-quality RNA for each sample was used for library preparation according to the Illumina Stranded mRNA Prep, Ligation (Illumina, 20040532). Library preparation and PolyA-RNA-seq on NovaSeq 6000 Sequencing System instrument (Illumina Inc., CA) was performed by the Genomics Facility, Italian Institute of Technology (IIT), Genoa, Italy. Sequencing was performed bidirectionally using a flow cell NovaSeq 6000 S1 Reagent Kit v1.5 (Illumina, 20028317) with the following parameters: 300 cycles (R1 length=150; R2 length=150), 40 million reads/sample. Small RNA sequencing: 1 µg of RNA for each sample was used for library preparation according to the NEXTFLEX Small RNA-Seq Kit v3 protocol (PerkinElmer, #NOVA-5132-06). Briefly, 3’ adapters were ligated to the 3’ end of RNAs using a 3’ RNA ligase enzyme followed by 5’ adaptor ligation using a 5’ RNA ligase enzyme. Reverse transcription followed by PCR was used to prepare cDNA using primers specific to the 3’ and 5’ adapters. The amplification of these fragments was carried out with 18 PCR cycles. The quality of the library was assessed by Bioanalyzer with Agilent High Sensitivity DNA kit (Agilent, 5067-4626). Small RNA-seq on NovaSeq 6000 Sequencing System instrument (Illumina Inc., CA) was performed by the Genomics Facility, Italian Institute of Technology (IIT), Genoa, Italy. Sequencing was performed bidirectionally using a flow cell NovaSeq 6000 SP Reagent Kit v1.5 (Illumina, 20028401) with the following parameters: 100 cycles (R1 length=50; R2 length=50), 20 million reads/sample.

RNA-seq analyses. Differential gene expression analysis was performed with QIAGEN CLC Genomic Workbench (QIAGEN CLC Genomics Server 25.0). Raw data were imported into CLC and mapped to the *Homo Sapiens* reference genome. Differential expression analysis was performed comparing the Pool-treated cells gene expression to the Scrambled RNA-treated cells, obtaining a list of differentially expressed genes (DEGs). This list was sorted by keeping only the statistically significant DEGs with a p-value<0.05. For small RNA-seq, Illumina row data were trimmed to remove the 3’ adapter using Cutadapt, with parameters –e 0.15, and tetramers removed from both ends. Likewise, the untrimmed reads were processed for tetramers and merged with the trimmed reads. Merged reads were then aligned with miRge3.0^6^. MiRge outputs were normalized to obtain relative TPM for each sample for the miRNA detected. TPM values for the 11 miRNAs of the Pool were selected, compared to their expression (TPM) in non-neoplastic human cerebellum samples (BioProject PRJNA752352 – accession number: GSE181520^7^), and plotted into a clustergram graph. Pathway analysis: The list of DEGs list was used to identify the DEGs involved in the positive regulation of cell migration (0030335), found on Gene Ontology and GO Annotations (<https://www.ebi.ac.uk/QuickGO/>).

**Proteomics and analysis.** Sample preparation: DAOY cells were cultured and transfected as described above at the final concentration of 10 nM. After 72h, proteins were extracted from 5 independent replicates per DAOY/per condition, prepared with ice-cold RIPA buffer containing protease inhibitors (Complete mini EDTA-free, Roche) and sodium orthovanadate (phosphatase inhibitor). Lysates were kept 30 min in agitation, centrifuged at 13000 x *g* for 30 min at 4°C and stored at -80°C until further processing. 50 ug of proteins in RIPA buffer were processed as follows: cysteine residues were reduced/alkylated with a final concentration of 10 mM TCEP and 30mM Chloroacetamide (95°C for 15 min in the dark). Samples were cooled at RT digested following the SP3 procedure^8^. The concentration of the resulting peptides was checked (Fluorometric Peptide Assay, ThermoFisher) and 300 ng of peptides were used for the proteomic analysis. The proteomics analysis was conducted on a Thermo Exploris 480 orbitrap system coupled with a Dionex Ultimate 3000 nano-LC system. After trapping and desalting, the tryptic peptides were loaded on an Aurora C18 column (75 mm x 250 mm, 1,6 µm particle size) nanocolumn (Ion Opticks, Fitzroy, Australia) and separated using a linear gradient of acetonitrile in water (both added with 0.1% formic acid), from 3% to 41% in 1 h, followed by column cleaning and reconditioning. The flow rate was set to 300 nL/min. Total run time was 1.5 h. Injection volume was set to 1 µL. LC/MS analysis: Peptides were analyzed in positive ESI mode, using a capillary voltage set to 2.0 kV. The RF lens was set to 40% and the AGC target was set to 300%. Data acquisition was performed in Data Independent mode (DIA) with a survey scan set from 400 to 1000 m/z at 120,000 resolution, followed by MS/MS acquisition of 60 m/z transmission windows, each having a fixed 10 Da width. MS/MS spectra were acquired in HCD mode. All the collected MS/MS spectra were analyzed using Spectronaut (Version 18), by running a DirectDIA analysis against the reference Homo Sapiens FASTA database (Tax ID: 9606 reporting 51548 reviewed entries). Cysteine carbamidomethylation was selected as fixed modification; acetylation of protein N-Term and Methionine oxidation were selected as variable modifications. Positive protein identifications were retained at 1% false discovery rate (FDR) threshold and at least two peptides were used for protein quantification. All the proteomics analysis were performed by the Analytical Chemistry Facility of the Istituto Italiano di Tecnologia (IIT), Genoa, Italy. Proteomics PCA analysis and graph generation were performed on custom MATLAB script (MathWorks, Natick MA).

**Statistical analyses.** Data were analyzed by GraphPad Prism 10 and are shown as mean ± Standard Error Mean (SEM). Statistical significance was evaluated with a two-tailed unpaired t-test for experiments in two groups at a single time point, two-tailed unpaired multiple t-test for experiments in two groups at a single time point for multiple independent elements, one-way ANOVA with Tukey’s multiple comparisons test, and two-way ANOVA with Bonferroni’s multiple comparison test for experiments with more than two groups and two variables. Transcriptomics statistics for long RNA-seq were performed through a Wald test. Results were considered significant for *p<0.05, **p<0.01, ***p<0.001, ****p<0.0001.

**SUPPLEMENTARY FIGURES (S1-S7)**

**Supplementary Figure S1**

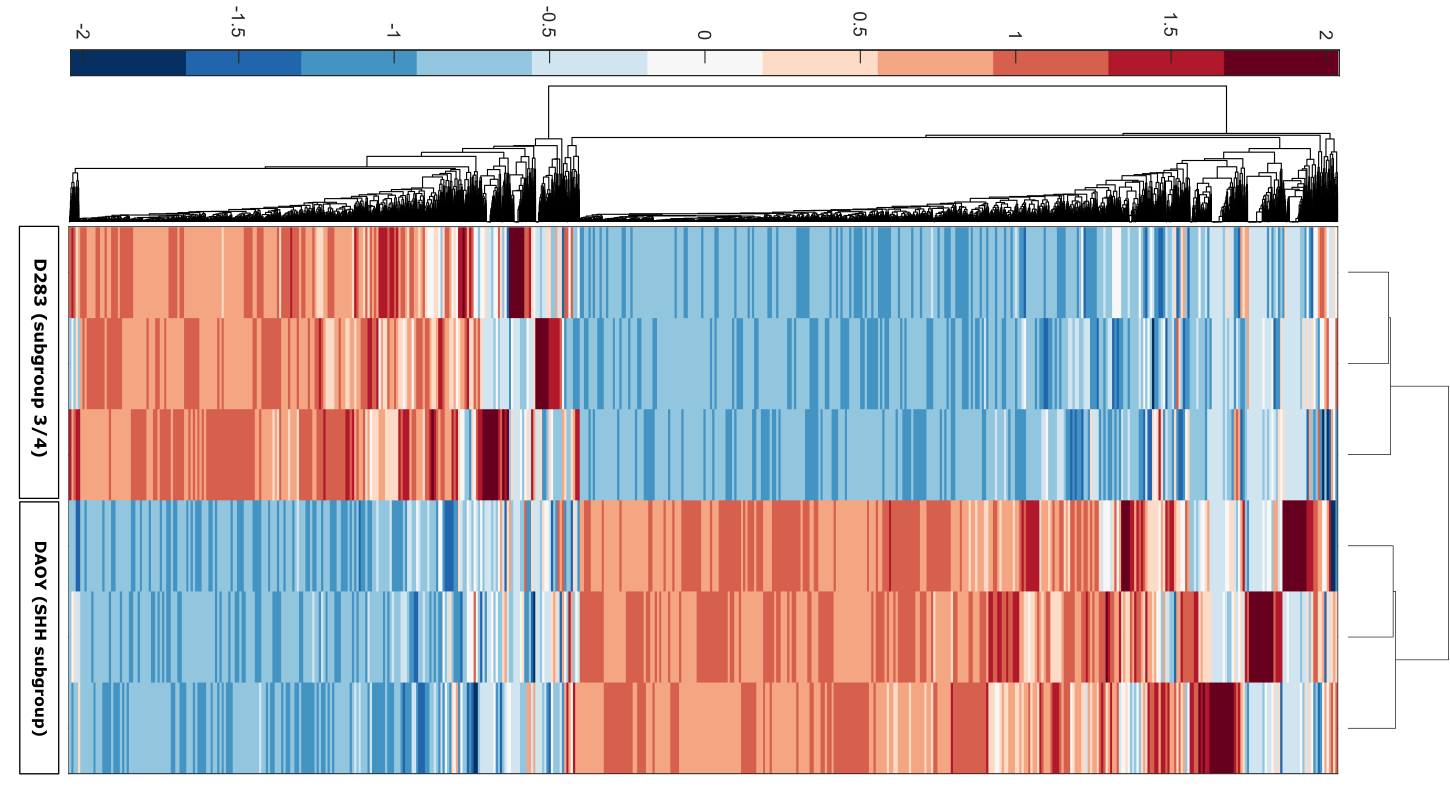

**Supplementary Figure S1. Differential genome expression in cell line models of different MB subgroups.** RNA-seq data of the gene expression landscape in D283 and DAOY cells. Data shown are the genes with an expression > 0 in both the cell lines. Samples were clustered for both genes and biological replicates.

**Supplementary Figure S2**

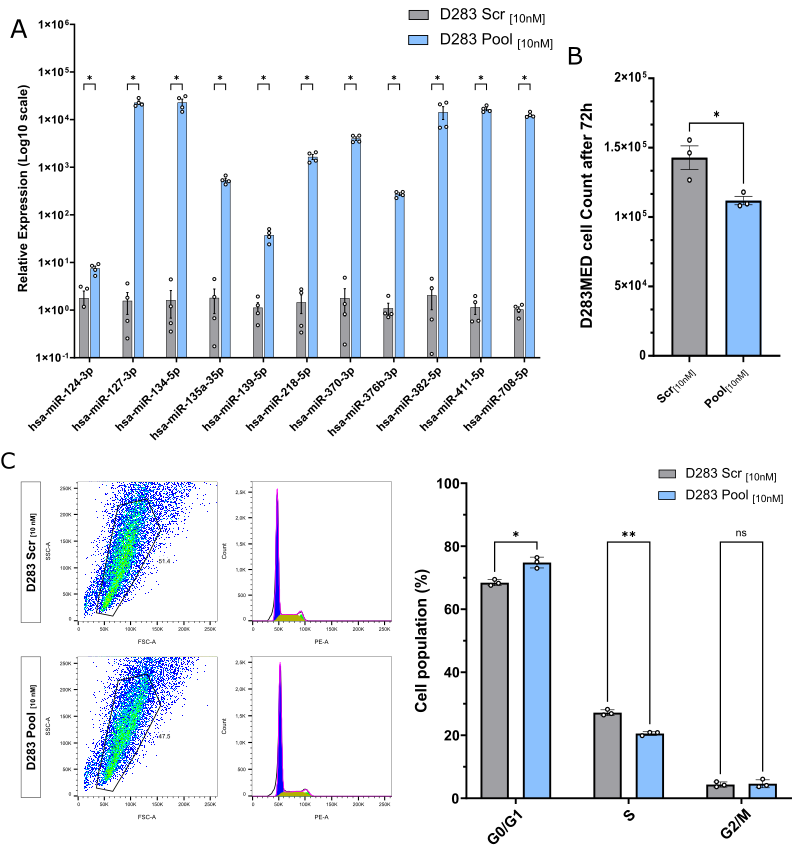

**Supplementary Figure S2. Restoration of the Pool in D283 inhibits cell growth**. (A) qPCR quantification of the indicated miRNAs upon transfection in D283 cells at the indicated concentration, compared to their levels in Scrambled-transfected cells (control). Statistical analyses were performed on n=4 biological replicates through a multiple t-test. (B) Bar plot indicating the cell count 72 hours after Pool or Scrambled RNA transfection at the indicated concentration. Statistical analyses were performed on n=3 biological replicates through a t-test. (C) Representative cell cycle analysis of PI staining by flow cytometry 72 hours after the transfection of D283 cells with the Pool or Scrambled RNA at the indicated concentration. Bar plots indicate the percentage of cells in the three cell cycle phases (G0/G1, S, G2/M). Data are represented as mean ± SEM of n=3 biological replicates. Multiple t-test., *p<0.05, **p<0.01, ***p<0.001.

**Supplementary Figure S3**

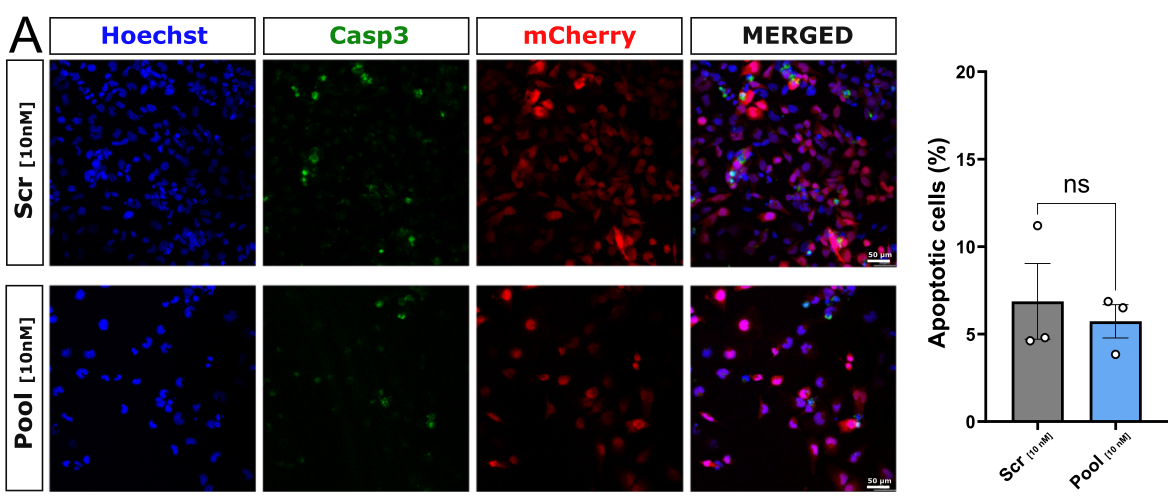

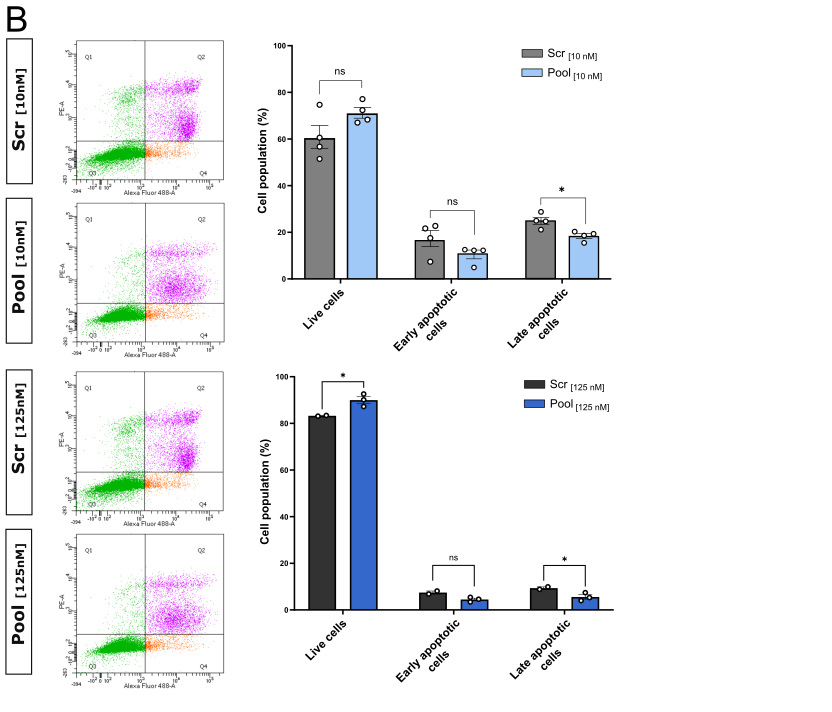

**Supplementary Figure S3 Transfection of the Pool does not induce apoptosis in DAOY cells.** (A) Immunofluorescence for cleaved-caspase3 in Pool- or Scramble-transfected DAOY cells after 72h. The apoptosis rate was calculated as Casp3^+^ cells over the total cell number. T-test on n=4 biological replicates. (B) FACS analysis of Annexin V-FITC^+^ cells (identified as early apoptotic – orange); Annexin V-FITC^+^/PI^+^ cells (identified as early apoptotic – pink) in Pool- or Scramble-transfected DAOY cells. Multiple t-test on n=3 biological replicates, *p<0.05, **p<0.01, ***p<0.001.

**Supplementary Figure S4**

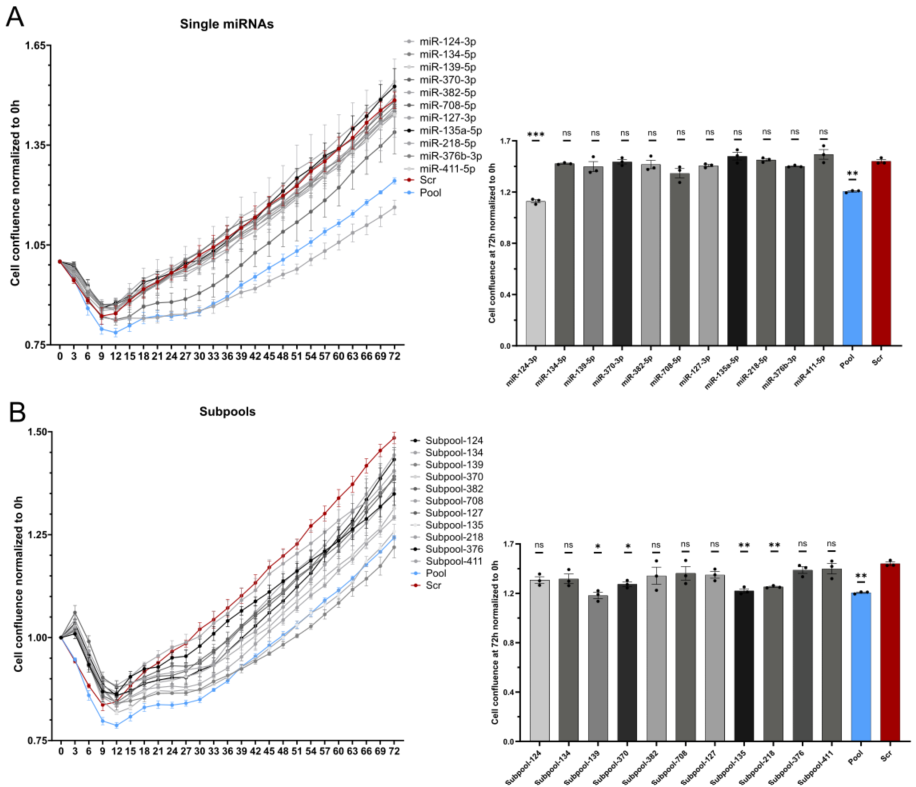

**Supplementary Figure S4. Synergistic effect of the Pool.** A) Quantification of DAOY cell growth over 72 hours (left) or cell confluence at 72h (right) upon the transfection of the Pool or Scrambled RNA, or individual miRNAs, and B) quantification of DAOY cell growth over 72 hours (left) or cell confluence at 72h (right) upon the transfection of the Pool, or Scrambled RNA, or subPools (Pool – miR*x*) at 10 nM. through live imaging-mediated of cell confluence over time. Two-way-ANOVA test (n=3). *p<0.05, **p<0.01, ***p<0.001.

**Supplementary Figure S5**

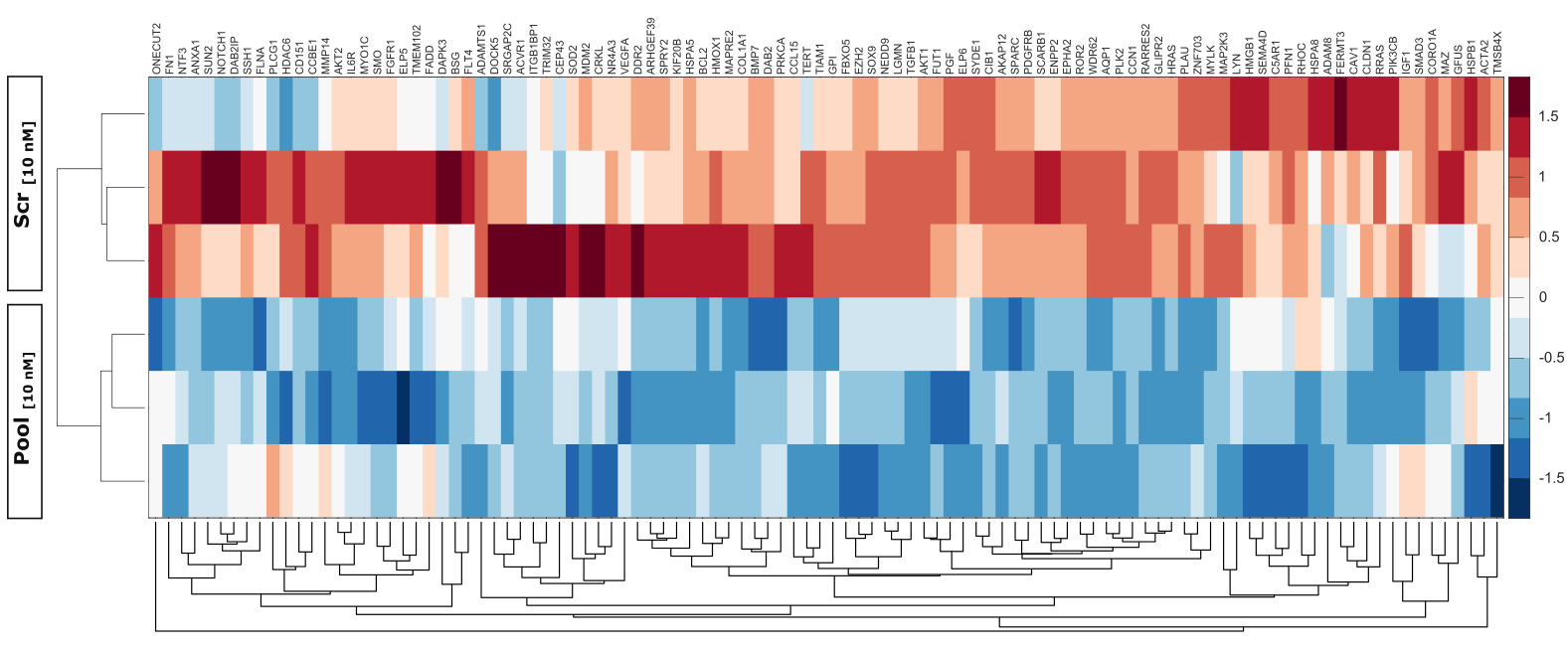

**Supplementary Figure S5. Restoration of the Pool in D283 inhibits expression of migration-related genes.** RNA-seq expression data of genes encoding for proteins involved in positive regulation of cell migration pathway (GO: 0030335) in Pool- or Scramble-transfected D283 cells. Samples were clustered based on biological samples (rows) and gene expression (columns). Only significantly downregulated genes are plotted. Statistical analyses were performed through Wald test on n= 3 biological replicates, *p<0.05, **p<0.01, ***p<0.001.

**Supplementary Figure S6**

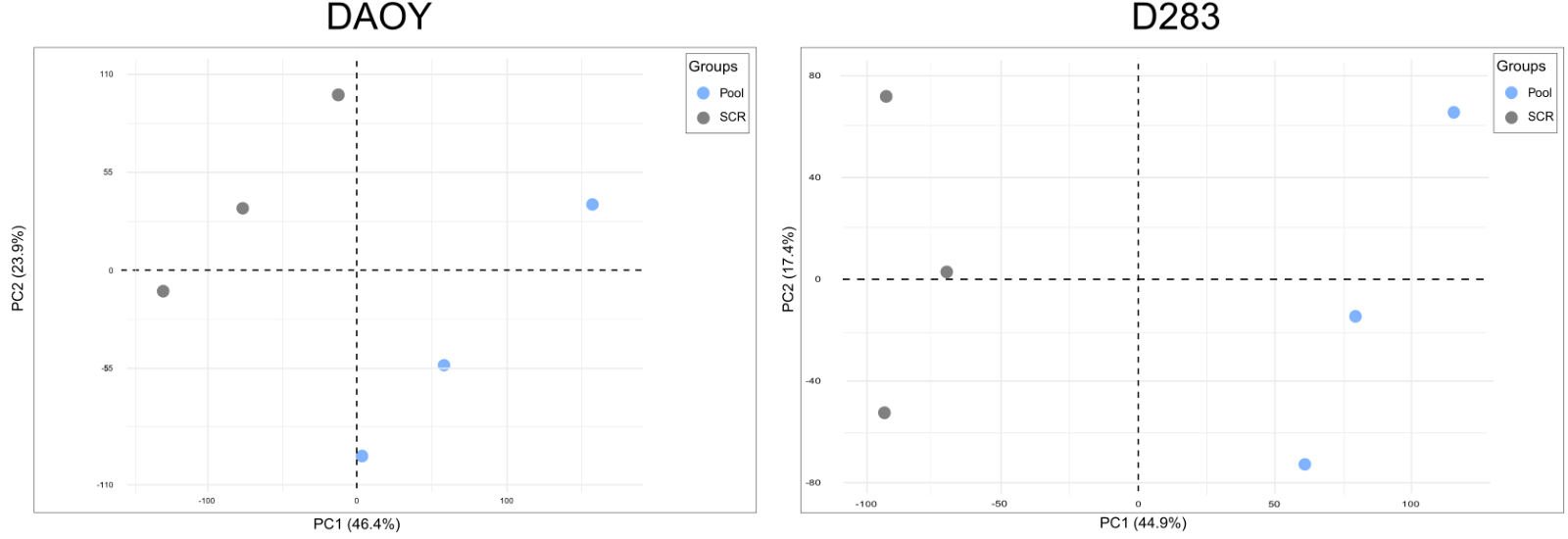

**Supplementary Figure S6. PCA** **of the transcripts** **in Pool- or Scrambled-transfected DAOY (*left*) and D283 (*right*)** (n=3 biological replicates).

**Supplementary Figure S7**

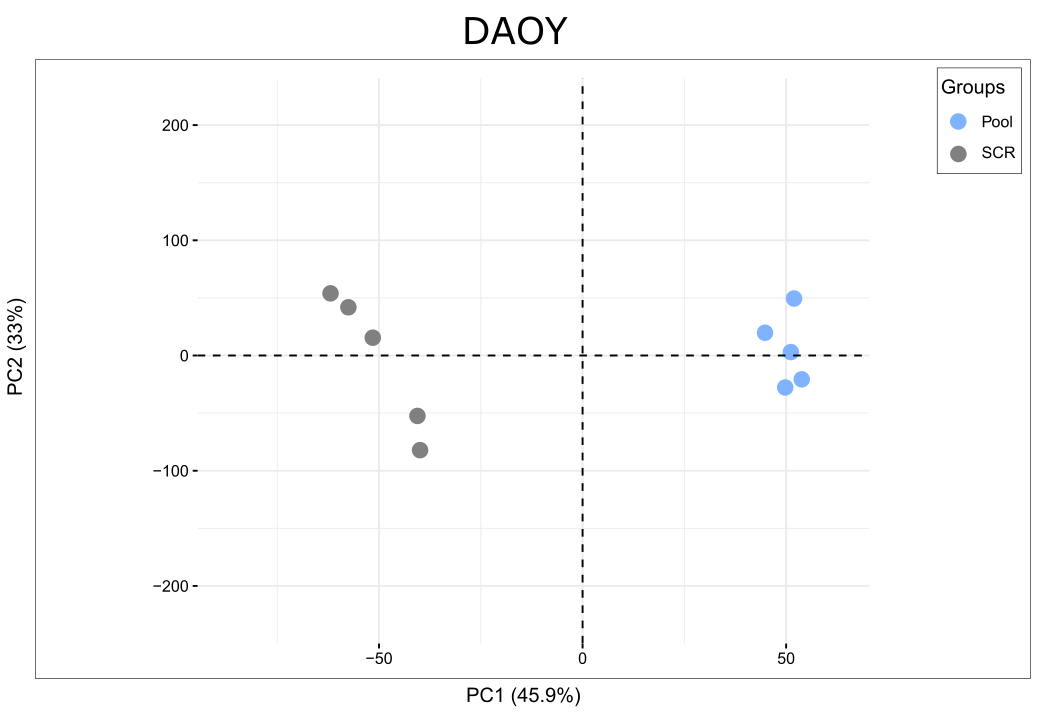

**Supplementary Figure S7**. **PCA of the proteins in Pool- or Scrambled- transfected DAOY** (n=5 biological replicates).

**SUPPLEMENTARY TABLES**

**Supplementary Table 1**

| miRNA name and accession number | miRNA sequence |
| --- | --- |
| mmu-miR-376b-3p MIMAT0001092 | AUCAUAGAGGAACAUCCACUU |
| hsa-miR-376a-3p MIMAT0000729 | AUCAUAGAGGAAAAUCCACGU |
| hsa-miR-376b-3p MIMAT0002172 | AUCAUAGAGGAAAAUCCAUGU |
| hsa-miR-376c-3p MIMAT000072 | AACAUAGAGGAAAUUCCACGU |

**Supplementary Table 1**. **Comparison among miR-376-3p sequences from mouse and human.** Mature miRNA sequences are written in 5’>3’ direction. Seed regions are underlined. Mismatches between mouse and human miRs are indicated in red.

**Supplementary Table 2**

| **miRIDIAN microRNA** | **Reference Dharmacon** | **Sequence** |
| --- | --- | --- |
| mmu-miR-135a-5p  hsa-miR-376b-3p  mmu-miR-139-5p  mmu-miR-218-5p  mmu-miR-411-5p  mmu-miR-127-3p  mmu-miR-134-5p  mmu-miR-370-3p  mmu-miR-382-5p  mmu-miR-708-5p  mmu-miR-124-3p | C-310411-05-0002  C-300741-03-0002  C-310568-07-0002  C-310576-05-0002  C-310822-01-0002  C-310397-07-0002  C-310409-05-0002  C-310619-07-0002  C-310607-05-0002  C-310987-01-0002  C-310391-05-0002 | UAAGGCACGCGGUGAAUGCC  AUCAUAGAGGAAAAUCCAUGU  UGUGACUGGUUGACCAGAGGGG  UAUGGCUUUUUAUUCCUAUGUGA  UCUACAGUGCACGUGUCUCCAG  UUGUGCUUGAUCUAACCAUGU  GCCUGCUGGGGUGGAACCUGGU  AUCAUAGAGGAACAUCCACUU  GAAGUUGUUCGUGGUGGAUUCG  UAGUAGACCGUAUAGCGUACG  AAGGAGCUUACAAUCUAGCUGGG |

**Supplementary Table 2. Sequences of the mature miRNA sequences and codes of the mimics.** Mature miRNAs mimics used for the *in vitro* transfection of the Pool. Except for human-specific hsa-miR-376b, all the other mimics are conserved between mouse and human.

**Supplementary Table 3**

| **Antigen** | **Vendor** | **Dilution** | **Antigen retrieval** |
| --- | --- | --- | --- |
| **Anti-BrdU** | Abcam (ab6362) | 1:100 | yes |
| **Anti-CD31** | BD pharmigen (550274) | 1:100 | no |
| **Anti-CD68** | Abcam (ab53444) | 1:100 | no |
| **Anti-mCherry** | Abcam (ab206402) | 1:500 | - |
| **Anti-Claudin-5** | ThermoFisher (4c3c2) | 1:100 | no |
| **Anti-IBA1** | Fujifilm (019-19741) | 1:500 | no |
| **Anti-Ki67** | Abcam (ab15580) | 1:100 | yes |
| **Anti-Laminin** | Abcam (7463) | 1:500 | no |
| **Anti-hNestin** | Abcam (ab105389) | 1:200 | no |
| **Anti-VEGF** | Abcam (ab52917) | 1:100 | no |
| **Anti- Cleaved Casp-3** | CellSignal (9664) | 1:800 | yes |

**Supplementary Table 3. List of primary antibodies used on immunofluorescence assays.**

**Supplementary Table 4**

| **Antigen** | **Vendor** | **Dilution** |
| --- | --- | --- |
| **Goat anti-rabbit AlexaFluor 647** | ThermoFisher (A21245) | 1:1000 |
| **Goat anti-rat AlexaFluor 488** | ThermoFisher (A11006) | 1:1000 |
| **Goat anti-chicken AlexaFluor 568** | Abcam (ab175477) | 1:1000 |
| **Goat anti-mouse AlexaFluor 488** | ThermoFisher (A11029) | 1:1000 |
| **Goat anti-rabbit AlexaFluor 647** | ThermoFisher (A21245) | 1:1000 |
| **Goat anti-rabbit AlexaFluor 568** | ThermoFisher (A11036) | 1:1000 |

**Supplementary Table 4. List of secondary antibodies used on immunofluorescence assays.**

**Supplementary Table 5**: **TPM values from long RNA-seq of DAOY** (*first sheet*). Transcriptomics results: significantly altered genes in Pool- vs Scrambled-transfected DAOY cells (10 nM, n=3 biological replicates) (*second sheet*).

**Supplementary Table 6**: **TPM values from long RNA-seq of D283** (*first sheet*). Significantly altered genes in Pool- vs Scrambled-transfected D283 cells (10 nM, n=3 biological replicates) (*second sheet*).

**Supplementary Table 7**: **Proteomics of DAOY.** Protein expression (*first sheet*) and significantly altered proteins (*second sheet*) in Pool- vs Scrambled-transfected DAOY cells (10 nM, n=5 biological replicates).
